# Principles for coding associative memories in a compact neural network

**DOI:** 10.1101/2020.06.20.162818

**Authors:** Chrisitian O. Pritz, Eyal Itskovits, Eduard Bokman, Rotem Ruach, Vladimir Gritsenko, Tal Nelken, Mai Menasherof, Aharon Azulay, Alon Zaslaver

**Affiliations:** Department of Genetics, Silberman Institute of Life Science, Edmond J. Safra Campus, The Hebrew University of Jerusalem, Jerusalem 9190401, Israel

## Abstract

A major goal in neuroscience is to elucidate the principles by which memories are stored in a neural network. Here, we have systematically studied how the four types of associative memories (short- and long-term memories, each formed using positive and negative associations) are encoded within the compact neural network of *C. elegans* worms. Interestingly, short-term, but not long-term, memories are evident in the sensory system. Long-term memories are relegated to inner layers of the network, allowing the sensory system to resume innate functionality. Furthermore, a small set of sensory neurons is allocated for coding short-term memories, a design that can increase memory capacity and limit non-innate behavioral responses. Notably, individual sensory neurons may code for the conditioned stimulus or the experience valence. Interneurons integrate these information to modulate animal behavior upon memory reactivation. This comprehensive study reveals basic principles by which memories are encoded within a neural network, and highlights the central roles of sensory neurons in memory formation.

## Introduction

Learning and memory processes are presumably universal in the animal kingdom, forming the basis for adaptive behavior. An intriguing form of these behavioral adaptations is known as associative learning, where a link between two unrelated cues is formed. The famous pavlovian dogs set a classical example: These dogs were trained to associate the sound of a bell (the conditioned stimulus, CS) with food (unconditioned stimulus, US). Consequently, the mere sound of the bell prompted the dogs to salivate in expectation for their meal (Pavlov, 1910).

To synthesize a productive associative memory that elicits an adaptive response upon future encounter with the CS, both the CS and the US must be encoded in the neural system. Moreover, their encoding needs to be logically integrated such that the behavioral response will match the expected valence that the CS predicts (Josselyn and Tonegawa, 2020). Whether the CS was associated with a positive or negative experience, this valence remains associated with the CS.

Animals have come up with different strategies for encoding associative memories. In flies, olfactory associative learning is centralized in the mushroom body, where both the valence of the US and the CS odorant are encoded (Widmann *et al.*, 2018). In contrast, mammalian brains are thought to encode associative memories in a decentralized fashion where interconnected areas, distributed throughout the entire brain, link up to encode memory traces. For example, associative fear memories are thought to be distributed among the amygdala that encodes the valence, the hippocampus which encodes the context, and the cortical neurons which provide the specific sensory information (Josselyn and Tonegawa, 2020).

Furthermore, sensory neurons proved to play intriguing roles in formation of associative memories. Their learning-induced neuroplasticity was observed across various sensory modalities (e.g. olfactory, gustatory, auditory, and visual), and is thought to confer improved detection and enhanced attention towards important cues encountered in the past (Åhs *et al.*, 2013; McGann, 2015).

Extracting the principles by which memories are formed within neural networks requires first to identify the brain regions, and ultimately the individual neurons, that participate in these processes. To this end, *Caenorhabditis elegans* worms offer an appealing research system. Their compact nervous system consists of 300 neurons, and a detailed blueprint of all the chemical and electrical connections is available (White *et al.*, 1986; Cook *et al.*, 2019; Witvliet *et al.*, 2020). Moreover, the number and the identity of the neurons is invariant and individual neurons can be unambiguously identified according to their position and anatomy across different individuals. Crucially, though equipped with a small neural network, *C. elegans* worms can form both associative and non-associative memories (Ardiel and Rankin, 2010; Sasakura and Mori, 2013). As in higher organisms, these memories can be classified into short- and long-term memories, which in worms last for a couple of hours or days, respectively (Kauffman *et al.*, 2010; Amano and Maruyama, 2011).

Whole-brain functional imaging enables measuring activity from large volumes in animals that express activity-dependent fluorescent markers (Ahrens *et al.*, 2013; Aimon *et al.*, 2019; Voleti *et al.*, 2019). In *C. elegans*, advanced microscopy techniques allow imaging neural dynamics of the entire network with cellular resolution in both stationary and freely-behaving animals (Schrödel *et al.*, 2013; Kato *et al.*, 2015; Nguyen *et al.*, 2016; Venkatachalam *et al.*, 2016; Toyoshima *et al.*, 2020).

Thus, studying neuroplasticity on the network-wide scale with cellular resolution allows addressing intriguing questions that were hitherto considered mainly based on theoretical grounds. For example, how does plasticity of individual sensory neurons code and integrate the stimulus and the valence of the experience? How many neural resources are required to form an associative memory, and more fundamentally, are there general principles by which associative memories are encoded within a neural network?

Here, we have systematically studied how the four types of associative memories (positive/negative associations, each formed as a short- or long-term memory) are encoded within the compact neural network of *C. elegans*. Short-term, but not long-term, memories are evident in the sensory system of the animal. Moreover, a limited set of sensory neurons codes each of the short-term memories where individual neurons may code the conditioned stimulus or the valence of the experience. The downstream interneuron layer codes both short- and long-term memories, and individual interneurons integrate both stimulus and valence information to dictate behavioral outputs.

## Results

We begin by establishing training paradigms that form robust traces of associative memories (Fig. 1A). Building on existing protocols (Bargmann, Hartwieg and Horvitz, 1993; Kauffman *et al.*, 2011), we trained *C. elegans* worms to form each of the four types of associative memories: short-term aversive (STAV), short-term appetitive (STAP), long-term aversive (LTAV), and long-term appetitive (LTAP). Notably, we used the same CS (the odorant butanone) for all training paradigms. This allowed us to extract memory traces that are unique to the training paradigm and that are independent of the specific CS used.

**Fig. 1.**
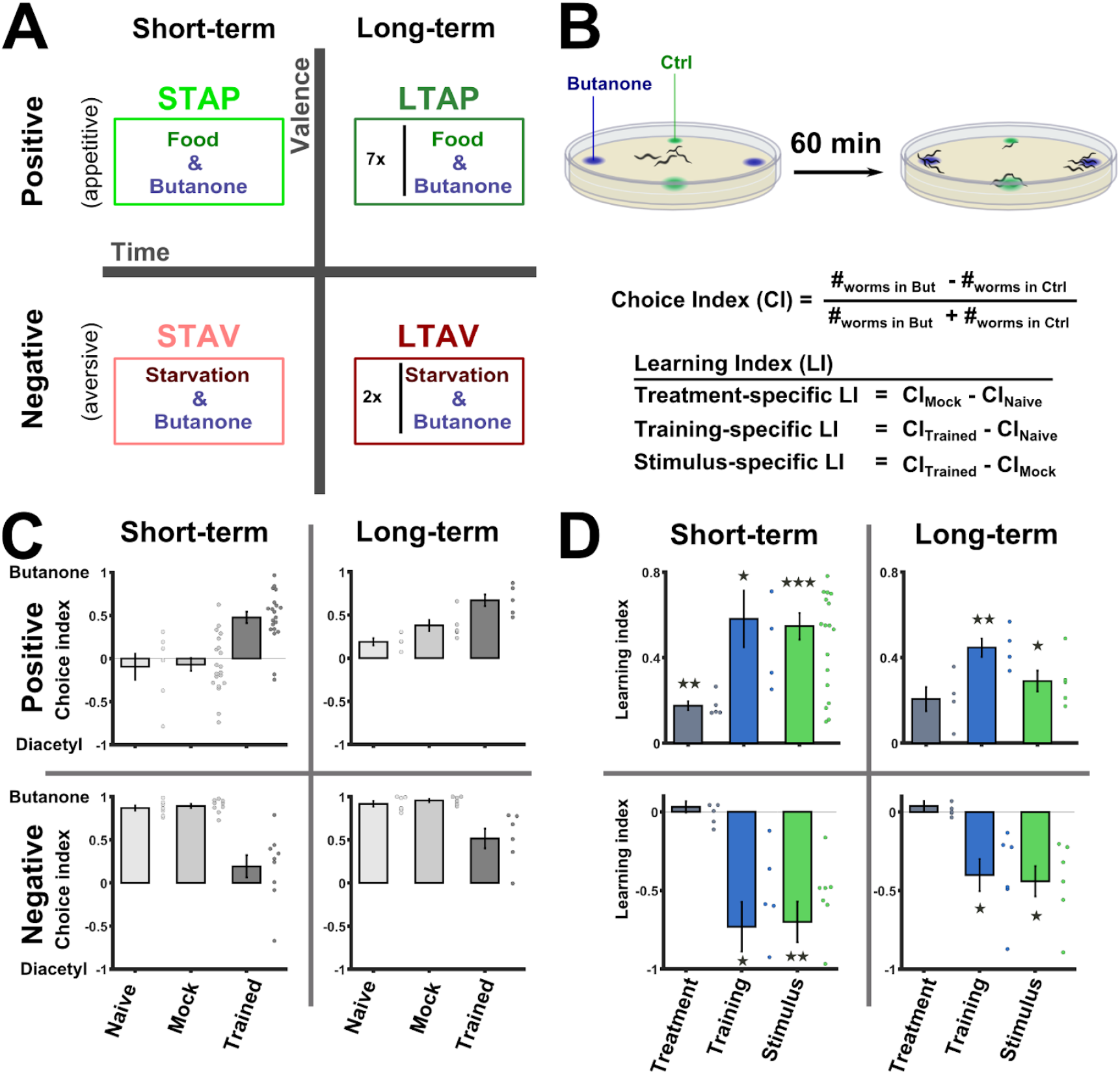
Training paradigms that form robust associative memories. **(A)** Worms were trained to form each of the four types of associative memories: Short- and long- term memories (denoted along the horizontal axis), each trained using a positive or negative unconditioned stimulus (vertical axis). Notably, the same conditioned stimulus, butanone, was used for all types of memory. STAP, short-term appetitive; LTAP, long-term appetitive; STAV, short-term aversive; LTAV, long-term aversive. **(B)** A two-choice assay was used to quantify animals’ preference towards the conditioned stimulus butanone (against the alternative attractive choice, diacetyl). Scoring the number of worms reaching each of the choices provided the Choice Index (CI), which ranges from −1 (denoting complete aversion to the CS) to +1 (full attraction). Choice tests for positively- and negatively- trained animals differed in concentrations and layout (see suppl. Fig. S1A) because of valence-specific effects on choice behavior (suppl. Fig. S2). Learning indices (LIs), calculated based on these CIs, show the treament-, stimulus-, and training- specific effects on the animals’ choice (suppl. Fig. S3). **(C)** Choice index values as scored following the behavioral choice assays. Positively-trained animals increased attraction while negatively-trained animals reduced attraction towards butanone. **(D)** Learning indices calculated according to the equations provided in (B) on the data shown in (C). Significant stimulus- and training-specific LIs in all paradigms indicate experience-dependent modulation of behavior that is based on stimulus and valence. Strikingly, Learning indices were calculated by comparing experiments performed on the same day. (suppl. Fig. S4 and Methods). Experimental repeats (C&D) were performed on different days and range between 4-21. Each experimental repeat is the average of three assay plates, each scoring 100-150 worms. Error bars indicate SEM. *p<0.05, **p<0.01, ***p<0.001 (one-sample t-test, FDR corrected; significant differences from the zero LI values).

To form positive (appetitive) or negative (aversive) associations, we exposed the worms to butanone in the presence or the absence of food, respectively. To form short-term memories, for which behavioral changes last for up to 2 hours (Kauffman *et al.*, 2010), we trained the worms for 1 hour and assayed the animals within 30 minutes following the training period. For long-term memories, which typically last for 1-2 days (>10% of the worms’ lifespan), we used repetitive-training regimes that lasted for ~12 hours and then assayed the animals 14 hours post the training period (Fig. 1A, Methods). In parallel to the trained animals, we always included mock-trained animals (animals that underwent training in the absence of the CS butanone) as well as naive animals, which were left untreated, as controls.

To verify that these training paradigms form robust memory traces, we analyzed the attraction of trained animals to the CS butanone (Fig. 1B). For this, we used a standard two-choice assay, where worms were free to choose between the CS and an alternative attractant, diacetyl, and calculated the Choice Index (CI), which provides a quantitative measure for the animals’ preference towards the CS (Fig. 1B, suppl. Fig. S1A).

Indeed, worms trained to form positive associations were significantly more attracted to butanone than mock-trained or naive animals (Fig. 1C-D). Similarly, worms trained to form negative associations, were significantly less attracted to butanone when compared to mock-trained or naive animals. These behavioral changes were evident in both short- and the long-term training paradigms. Similar results were also obtained when using ethanol (butanone diluent) instead of diacetyl as the alternative choice (suppl. Fig. S1B).

We next quantified the explicit effects of the CS (butanone) and the US (starvation/appetitive experience) on the behavioral output. For this, we used the CI values to compute the Learning Index (LI), the difference between the choice indices of the different experimental groups within a training paradigm (Fig. 1B and Methods). The treatment itself (CI_mock_ - CI_naive_) had negligible effects, except for a mild effect noted following the STAP- training protocol (Fig 1D). However, in all paradigms, significant changes in choice (both training (CI_trained_ - CI_naive_) and stimulus (CI_trained_ - CI_mock_) -specific) indicated that the modulated choice of butanone was due to the presence of butanone during training. Furthermore, the change in the choice corresponded to the valence of the experience: appetitively-trained animals increased their choice of butanone, while aversively-trained animals decreased their choice. Thus, the change in choice behavior is dependent on both the stimulus and the valence of the experience.

Taken together, using the same CS butanone, we could robustly form all the four types of associative memories: positive (appetitive) associative memories which increased attraction towards butanone, and negative (starvation) associative memories which decreased this attraction. Both negative and positive experiences could be used to form short- and long-term memories.

Memory formation is typically associated with changes in synaptic strength among neurons that participate in encoding the memory (Byrne, Castellucci and Kandel, 1978; Hopfield, 1982). As a result, activity of these memory-storing neurons may be modulated when compared to their activity in naive or mock-trained animals. In that respect, quantifying calcium dynamics within individual neurons is a powerful technique to extract fine neural dynamics (Kerr *et al.*, 2000; Ahrens *et al.*, 2013).

We begin by analyzing neuroplasticity of the sensory system, naturally focusing on the chemosensory neurons. To measure response dynamics in individual chemosensory neurons, we imaged a transgenic strain that expresses the genetically encoded calcium indicator, GCaMP, in virtually all chemosensory neurons (Iwanir *et al.*, 2019). We restrained the animals in a microfluidic device (Chronis, Zimmer and Bargmann, 2007), and used a fast-scanning confocal system to collect fluorescent image stacks of all chemosensory neurons during exposure to the CS butanone (Fig. 2A).

**Figure 2.**
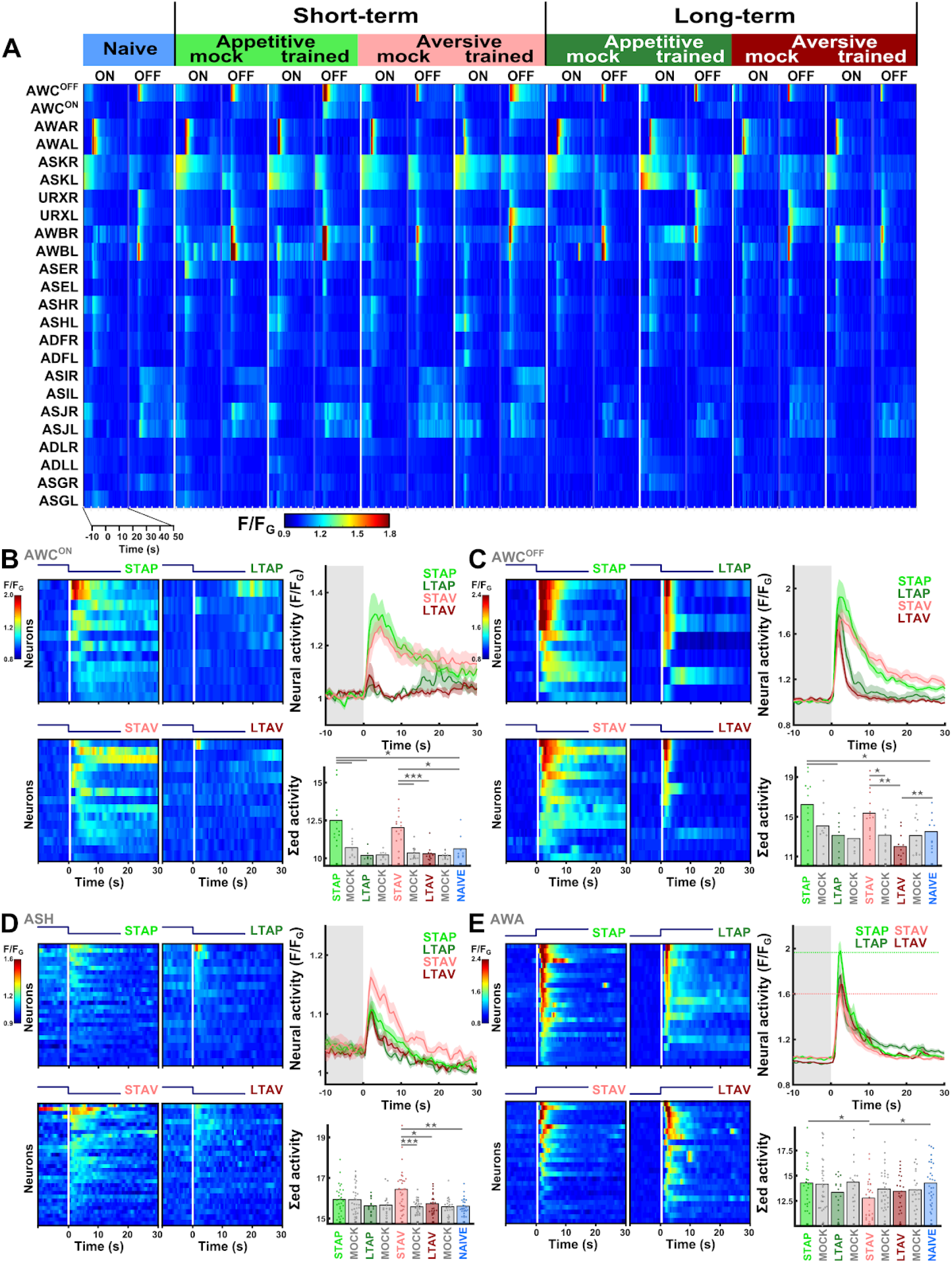
Short term, but not long-term, memories are evident at the sensory layer. **(A)** Neural dynamics following exposure to (ON) or removal of (OFF) the conditioned stimulus butanone. These activity dynamics were systematically measured in animals trained in the four associative-memory paradigms (STAP, STAV, LTAP, LTAV, Fig. 1A), as well as in their matched mock-trained and naive control animals. 9-17 animals were imaged per group (column), resulting in coverage of 2-17 per neuronal trace (median=13). **(B-E)** The sensory neurons, AWC^ON^ (B), AWC^OFF^ (C), ASH (D), and AWA (E) show significantly differential activity following formation of short-term memories. Heat maps denote activities of individual neurons in each of the training paradigms. Horizontal white line at t=0 denotes the time of stimulus exchange. In the mean line plots, the different colors denote the trained animals in a given paradigm. Shaded gray background indicates butanone exposure, and shaded area around the mean activity indicates standard error of mean. Bar graphs represent the summed neuronal activity post the stimulus switch. Blue, naive worms; Gray, mock-trained worms. Next to each bar, the groups that show significant differences are indicated. *p<0.05, **p<0.01, ***p<0.001 (t-test, FDR corrected). **(B)** In the AWC^ON^ neuron, STAV- and STAP-trained animals showed similar elevated responses, which were significantly higher than their matched mock-trained controls (bar graphs and suppl. Fig. S8), suggesting stimulus-specific encoding. **(C)** In the AWC^OFF^ neuron, responses following STAV- and STAP-training were significnatly increased. STAV-trained animals exhibited significantly higher responses when compared to the mock-trained animals (bar graphs and suppl. Fig. S8), suggesting stimulus-specific coding. **(D)** In ASH neurons, activity is increased after STAV training only. Note the significant difference from the mock-trained controls. ASHL and ASHR showed similar response dynamics, so shown are their averaged activities. **(E)** STAV-trained animals showed significantly higher responses than STAP-trained animals in AWA neurons, suggesting valence-specific differences. Horizontal dotted lines in line graphs highlight amplitude differences between STAV and STAP. AWAL and AWAR showed similar response dynamics, so presented are their averaged activities.

Naive *C. elegans* worms showed robust innate responses in AWA, ASH, AWC, AWB, ASJ, and URX sensory neurons towards exposure or removal of butanone (Fig. 2A and suppl. Fig. S5, S6 and in agreement with previous reports (Bargmann, Hartwieg and Horvitz, 1993;Kato *et al.*, 2014). However, following training and memory formation, response activity of merely four neuron types, namely, AWC, ASH, AWA and ASI, was markedly modulated (Fig. 2B-G,E, suppl. Figs. S7, S8). These neural modulations were observed only in the short-term memory paradigms, and none of these changes persisted in the long-lasting memories (see suppl. Fig. S9, except for the ASI neurons, whose modulated dynamics appeared only in half of the population, suppl. Fig. S7).

The two bilateral, symmetric AWC^ON^ and AWC^OFF^ neurons showed increased activity only in the short-term paradigms (Fig. 2B-C, suppl. Fig. S8 A-D). While the AWC^OFF^ neuron exhibited robust innate responses in naive animals, its responses were significantly elevated following formation of short-term memories (suppl. Fig. S10). The AWC^ON^ neuron lacked activity responses in naive and mock-trained animals, but gained marked responses following formation of short-term memories (suppl. Fig. S10). Notably, these observations were robustly replicated in an additional and independent strain that expresses GCaMP exclusively in the AWC^ON^ and AWC^OFF^ neurons, and which allowed unambiguously identifying each of these neurons (suppl. Fig. S10 and see Methods). Moreover, the gained responses in STAP-trained animals were dependent on intact synaptic activity (but not on neuropeptide signaling, suppl. fig. S11), indicating that active network dynamics is required for modulating AWC^ON^ responses.

For both AWC neurons, the response dynamics of positively- and the negatively-trained animals did not significantly differ (suppl. Fig. S10 A&B), suggesting that these neurons do not distinguish between the valence (the appetitive or aversive experience) of the memory. In contrast, the responses of the trained animals were significantly higher than their matched mock-trained controls (which were not exposed to the CS, Fig. 2 B & C bar graphs, suppl. Fig. S10 A&B), suggesting that these neurons may be coding the conditioned stimulus (suppl. fig. S3), a finding that is in line with previous reports (Cho *et al.*, 2016).

The ASH-type neurons showed enhanced responses following short-term negative training (Fig. 2D & suppl. Fig. 8C). These responses were significantly stronger than the responses of the matched mock-control animals, suggesting that the ASH neurons may be coding the stimulus. However, since the increased response was observed following the negative training only (Fig. 2D), it is also possible that these neurons code for the valence component of the memory.

The AWA-type neurons showed mild, yet significant, differential responses between the positive and the negative short-training paradigms (Fig. 2E). These responses were not different from their matched mock-trained controls, suggesting that the AWA neurons may be coding the experience (valence) rather than the stimulus (Fig. 2E, suppl. Fig. S3).

To further assess the role of these neurons in formation of short-term memories, we quantified the preference of trained animals, defective in each of these individual neuron types, towards the CS (as shown in Fig. 1A). Animals with genetically-ablated AWC neurons lost their learning and memory abilities as indicated by the near zero LI values for both the positive and the negative training paradigms (suppl. Fig. 12 A-B). Animals with functionally-defective AWA neurons (*odr-7*^-/-^), showed impaired learning abilities following negative training only, while animals with genetically-ablated ASH neurons were not significantly imapired (suppl. Fig. 12 A-B). These results indicate that neurons showing modulated activity in memory retrieval are indeed critical for memory formation or memory retrieval (or both) processes.

Together, these findings demonstrate that short-term, but not long-term, memories are encoded within a limited set of sensory neurons. Moreover, individual sensory neurons may code for either the stimulus or the valence (experience), or possibly like the ASH neurons, integrate the stimulus and the valence information.

As activity of sensory neurons was not modulated following formation of long-term memories (both LTAP and LTAV, fig. 2A and suppl. Fig. S9), we speculated that long-term memory traces may have shifted to the downstream interneuron layer. Since short-term memories showed modulated activity primarily in AWC, ASH and AWA neurons (Fig. 2), we studied their main shared downstream interneurons, namely, AIA, RIA and AIB (Fig. 3, suppl. Fig. S13 and S14).

**Figure 3.**
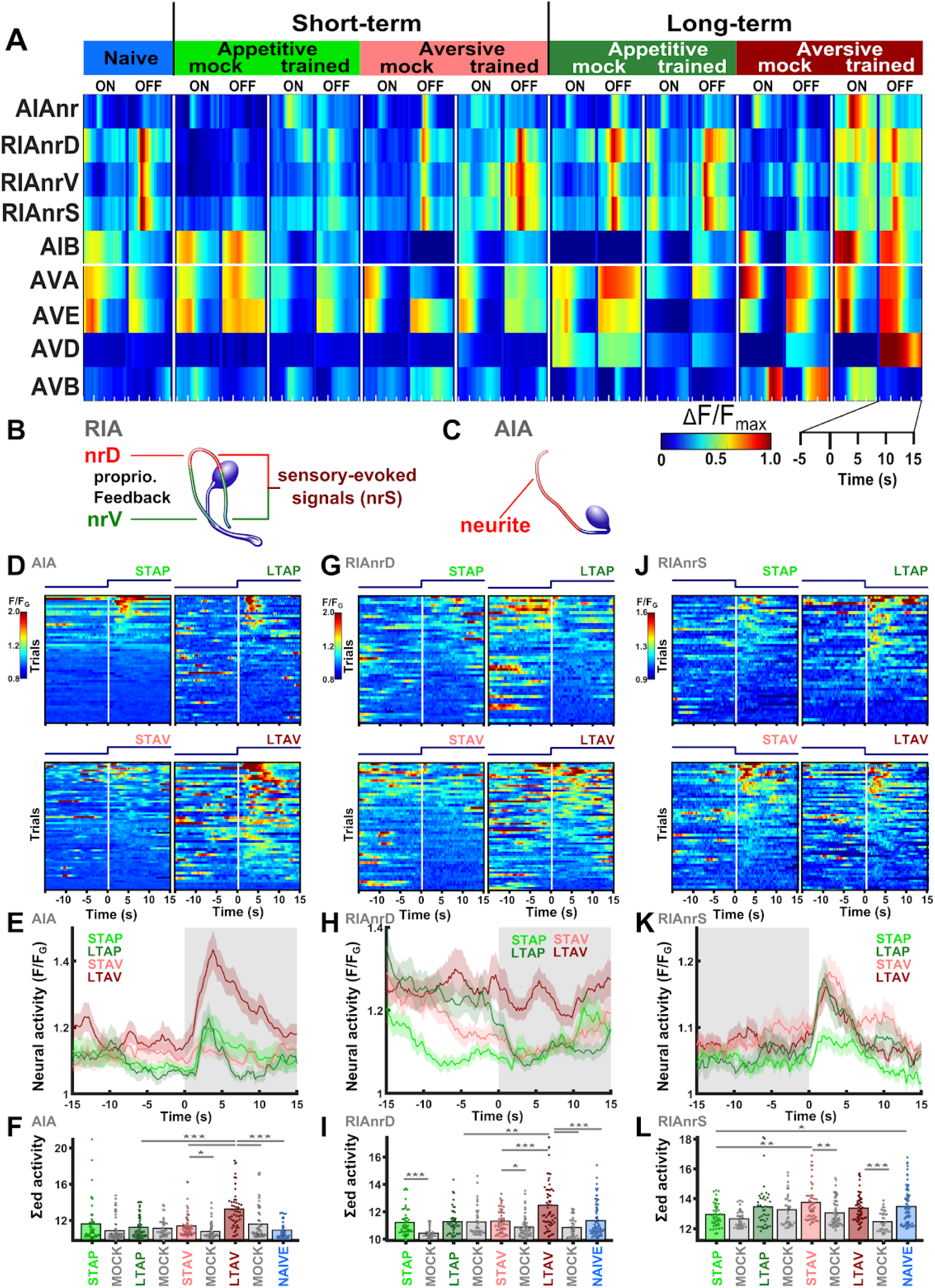
Interneurons integrate valence and stimulus information of short- and long- term memories. **(A)** Activity profiles of inter- and command- neurons following memory formation according to each of the four training paradigms (STAP, STAV, LTAP, LTAV, Fig. 1A). Butanone presentation (ON) or removal (OFF) was at t=0. ‘nr’ next to the neuron name denotes imaging Ca dynamics from the neurite. D, dorsal; V, ventral; S, stimulus-evoked signal. Each neural trace is the mean of at least four repeats. 9-15 individual animals were imaged. **(B)** Activity of the RIA neurons was measured in the dorsal (RIAnrD, red) and the ventral (RIAnrV, green) compartments of the neurite. The sensory-evoked signal (RIAnrS) was extracted as the overlap in signal from both the dorsal and the ventral compartments upon stimulus removal. **(C)** Activity in the AIA neurons was measured from the neurite (red region). **(D)** Heat maps, **(E)** line plots, and **(F)** bar graphs depicting activity in the AIA neurite (AIAnr). Note the increased activity following LTAV training and the differential activity following LTAP and associated mock-trained controls (stimulus- and valence specific differences). **(G)** Heat maps, **(H)** line plots, and **(I)** bar graphs depicting activity in the RIA dorsal neurite (RIAnrD). LTAV-trained animals show differential activity when compared to other training paradigms or their matched mock-trained controls, suggesting for stimulus and valence-specific differences (see also line graph & suppl. Fig. S13). **(J)** Heatmaps, **(K)** line plots, and **(L)** bar graphs depicting activity in the RIA dorsal neurite (RIAnrD). Note the increased activity in STAV- compared to STAP- training and mock-trained controls (stimulus and valence-specific differences, also see suppl Fig. S13). Line graphs: colors denote the trained animals in each of the training paradigms. Shaded areas in the line plots indicate SEM. Bar graphs represent the summed neuronal activity post stimulus exchange. * p<0.05, ** p<0.01, *** p<0.001, t-test, FDR corrected.

For this, we repeated the four training paradigms in strains expressing GCaMP in these interneurons. While all three neurons showed innate responses to butanone in naive animals, both AIA and RIA (but not AIB) showed significantly altered responses to butanone following memory formation (Fig. 3). As previously reported for AIA and RIA (Chalasani *et al.*, 2010; Hendricks *et al.*, 2012), these modulated dynamics were observed in the neurites rather than in the cell soma (Fig 3 B & C).

Following LTAV-training, AIA activity was significantly elevated compared to both naive and mock-trained controls (Fig. 3D-F), suggesting that the AIA neurons may be coding the conditioned stimulus. The enhanced response in LTAV-trained animals was also significantly higher than the response found in LTAP-trained animals, suggesting that these neurons may also code the negative experience. Thus, AIA neurons may be integrating information of both the stimulus and the valence.

The RIA neurons showed an interesting modulated dynamics. These neurons integrate proprioceptive feedback, that indicates the animal’s head position, together with sensory inputs (Hendricks *et al.*, 2012; Ouellette *et al.*, 2018). Remarkably, following long-term training paradigms, the baseline activity of the RIA neurons was elevated before the exposure to the stimulus (compare heat maps from Fig. 3G to heat maps in suppl. Fig. S13B). Following stimulus presentation, only in LTAV-trained animals activity in the dorsal neurite was maintained at high levels, while in all other training conditions activity markedly decreased (Fig. 3G-I). Similar, though less pronounced, results were obtained by imaging the ventral neurite of the neuron (suppl. Fig. S13E & G), where LTAV-trained animals showed highest post-stimulus activity. The differential activity between LTAV and LTAP training, and the fact that LTAV-trained animals showed higher activity than mock-control animals (Fig. 3F), suggest that the RIA neurons may encode the valence and the stimulus components of the memory.

We next extracted RIA activities that stem from the sensory input alone (Hendricks *et al.*, 2012; Ouellette *et al.*, 2018). These inputs were evident in both the ventral and the dorsal compartments of the neurite following stimulus removal (Fig. 3B, J, K, & L). However, the sensory-evoked activity was significantly prolonged in STAV-trained animals (compared to STAP- and mock- trained animals, Fig 3F and suppl Fig. S13D). This suggests that, in addition to long-term memories, the RIA interneurons integrate stimulus- and valence- sensory information stored in short-term memory. As RIA neurons directly connect to motor outputs, integrated information of stored and newly-sensed inputs can be efficiently mediated to respective behavioral outputs.

Command neurons, which directly control locomotive outputs, form a functional layer downstream of the above studied interneurons (Chalfie, 1984). We therefore studied whether activity dynamics in command neurons is also modulated following memory acquisition. For this, we used a transgenic strain that expresses GCaMP pan-neuronally (Nguyen et al 2017), from which we could reliably extract activities of the backward-motion promoting neurons AVA, AVE, AVD as well as from the forward-motion promoting neuron AVB (Fig. 3A and suppl. Fig. S14). Indeed, activities among the backward command neurons were highly correlated, and anti-correlated to the activity of the forward command neuron (suppl. Fig. S14). Based on the activity dynamics of the AVA neurons, whose response was most prominent, we could not detect differences in the response dynamics following the various training paradigms (suppl. Fig. S14B-E), suggesting that these neurons do not participate (or at the very least, play a minor role) in memory encoding.

Together, these findings demonstrate that activity of individual interneurons is modulated by the valence and the stimulus in both short- and long-term memories. Thus, individual interneurons may serve as integrating hubs for the encoded memory information.

The comprehensive analysis of neural activity showed that learning modulates activity dynamics of sensory- and inter- neurons (Figs. 2-3), which in turn, modulates animal attraction towards the training stimulus (Fig. 1). We next asked what are the fine locomotive changes that follow memory acquisition and which underlie the modulated attraction towards the CS. For this, we imaged trained worms during chemotaxis towards butanone (Fig. 4A) using a multi-animal tracking system that extracts key locomotion parameters from over a hundred worms at a time (Itskovits *et al.*, 2017). As expected, aversively-trained animals were significantly slower to arrive at the butanone target point when compared to positively-trained or naive animals (Fig. 4B, consistent with the chemotaxis results shown in Fig. 1C). However, positive training yielded no improvement as both trained and mock-trained controls were slower to leave the starting point area, presumably due to the specific technicalities of the assay (see suppl. Fig. S15C).

**Figure 4.**
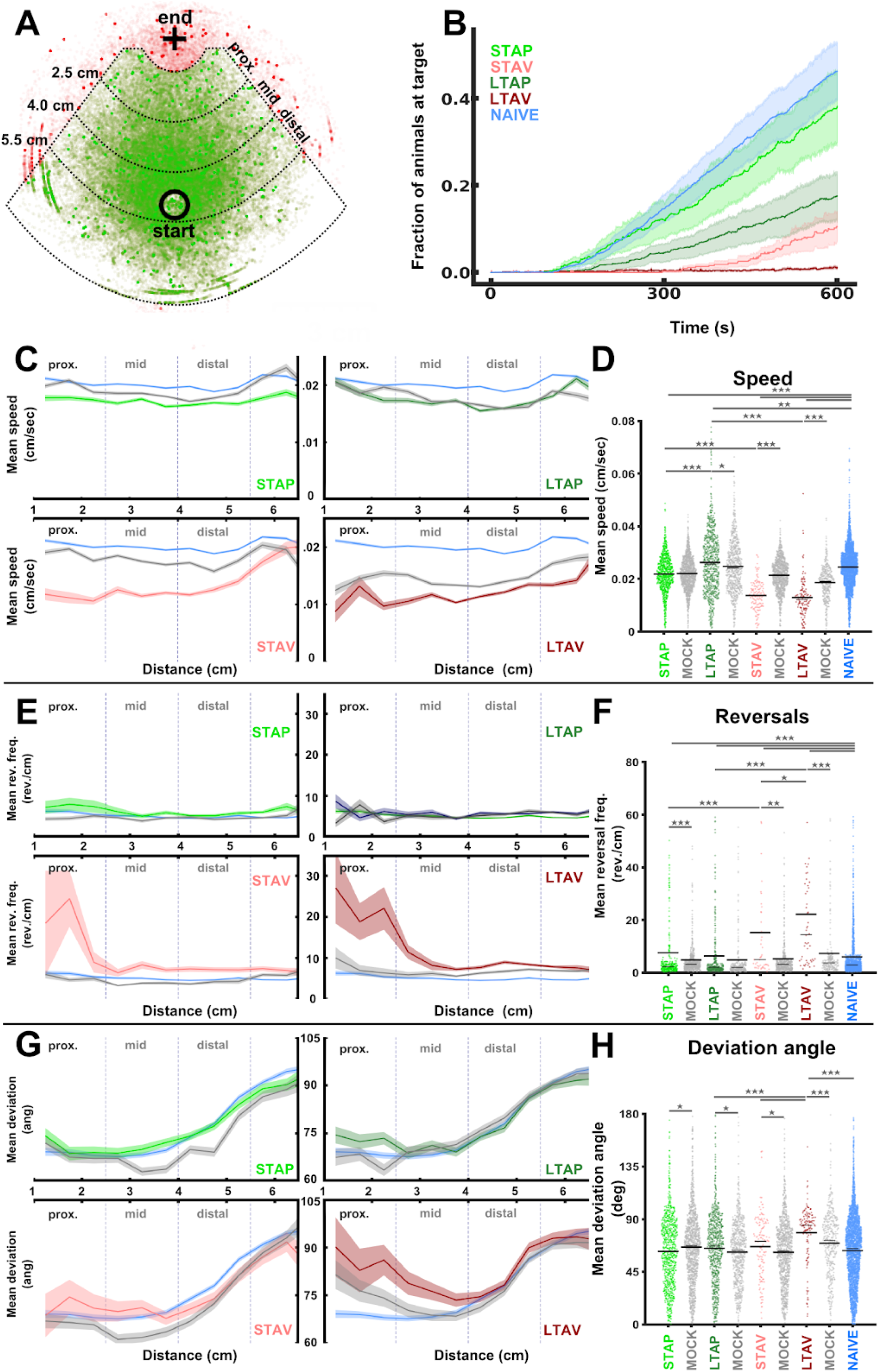
Memory-induced neural changes modify fine behavioral outputs. **(A)** layout of the one-choice chemotaxis assay. Butanone was applied on a chemotaxis plate (marked as + region), and ~100 worms, placed 5 cm away (black circle), were imaged during 10 minutes of chemotaxis. For analysis purposes, the arena was divided into three radial segments: proximal, mid., and distal, spaced 1.5 cm apart. Tracks bordering the plate rim were excluded (red points). All other tracks (green) were collected to extract locomotion parameters, as shown in panels C-H. **(B)** Temporal accumulation of animals at the butanone target region (marked + in panel A). Positively-trained and naive animals reached the target point faster than aversively-trained animals. 4-11 independent experimental repeats for each of the different conditions (21 for naive). Shaded areas denote standard error (estimated by bootstrap analysis). **(C)** Mean locomotion speed as a function of the distance from the target point. **(D)** Mean locomotion speed in the proximal zone. Note that speed is significantly reduced in aversively trained animals. **(E)** Mean reversal rates as a function of the distance from the target point for each of the four training paradigms. **(F)** Mean reversal rates in the proximal zone only. Reversals are significantly increased in aversively trained animals. **(G)** Deviation angles of animal’s trajectories from the target CS point. The deviation angles are plotted as a function of the animal’s distance from the target for each of the four training paradigms. **(H)** Deviation angles in the proximal zone only. Note that the deviation angle is significantly increased in LTAV-trained animals. In D,F,H: Asterisks and notations indicate significant differences. Error bars indicate SEM. *p<0.05, **p<0.01, ***p<0.001 (rank sum test, FDR corrected).

To extract the mechanisms underlying the diminished attraction towards butanone following aversive training, we analyzed key chemotaxis properties, including speed, turning frequency, and the directionality of the animals in relation to the CS endpoint (Fig. 4C-H and suppl. Fig. S15). Notably, negatively-trained animals were significantly slower during chemotaxis (compared to naive, mock-trained and positively-trained worms, Fig. 4 C,D and suppl. Fig. S15D-F). Moreover, the reduced speed was irrespective of the distance from the CS target point.

Strikingly, negatively-trained, and particularly LTAV-trained animals, showed increased reversal rates when reaching the proximity of the CS (compared to naive, mock-trained controls, and positively-trained animals, Fig. 4E&F, suppl. Fig. S15G). Furthermore, the trajectories of LTAV-trained animals became less directed towards the target CS as reflected by their increased deviation angle (Fig. 4G&H, suppl. Fig. S15H), indicating their aversion once approaching close to the CS.

Thus, formation of negative associations lead to marked changes in key locomotion parameters, including speed, turning frequency, and directionality towards the target containing the aversive conditioned stimulus. The combined effect of these behavioral modulations can explain the diminished attraction towards the CS following formation of negative associations.

## Discussion

Here, we have systematically studied how associative memories are encoded within the compact neural network of *C. elegans*. Specifically, we elucidated the coding scheme for each of the four forms of associative memories: short- and long- term memories, each coded using a positive and a negative association. Importantly, the use of the same conditioned stimulus (CS, butanone) in all memory paradigms allowed us to extract the individual neurons that exclusively code the experience valence (positive or negative), the conditioned stimulus, or both.

Our findings indicate that short-term, but not long-term, memories modulate response dynamics of sensory neurons towards the CS (Fig. 2). We observed significant changes in at least three sensory neuron types following formation of short-term memories. The AWC neurons showed clear modulated responses in a stimulus-specific manner (Fig. 2B-C), in line with previous findings showing that the AWC neurons represent sensory history by shifting their susceptibility towards the trained concentration of the CS (Cho *et al.*, 2016). Interestingly, the modulated activity in the AWC^ON^ neuron depended on intact neurotransmission (Suppl. Fig. S11), demonstrating that network activity, rather than cell-autonomous processes, underlie memory formation in this neuron.

In addition to coding sensory history, we found that sensory neurons may also code the valence of the experience. This was observed in the AWA neurons, and possibly also in the ASH neurons (Fig. 2D-E). Thus, valence and stimulus components of the memory may be distributed among individual sensory neurons, and integration of their modulated dynamics, presumably, forms the associative memories. A division of encoding valence and the stimulus components of the memory is also found in insects and mammalian brains (Gottfried *et al.*, 2002; Sacco and Sacchetti, 2010; Sekeres *et al.*, 2010; Liu *et al.*, 2012). It appears that modularization of memory components is a fundamental principle that even compact nervous systems adhere to.

These findings highlight the intriguing perspective that sensory systems are actively participating to encode associative memories. In mammals, similar stimulus-specific neuroplasticity was observed in primary sensory cortices (Morris, Friston and Dolan, 1998; Ohl and Scheich, 2005) and in peripheral sensory neurons (Jones *et al.*, 2008). Thus, neuroplasticity at the sensory system may have evolved as a universal solution to improve detection and to enhance attention towards critical stimuli, thereby increasing animals’ fitness (McGann, 2015).

A limited set of chemosensory neurons was recruited to code each of the short-term memories. Out of the 12 pairs of sensory neurons analyzed herein, only a single pair (AWC) showed modulated activity following short-term positive training, and three pairs of neurons (AWC, ASH, AWA) showed modulated activity following short-term aversive training (Fig. 2). This activity sparseness may come as a surprise given the considerable connectivity among the sensory neurons, either by direct wiring (chemical and electrical) or via feedback from connected interneurons (White *et al.*, 1986; Zaslaver *et al.*, 2015; Cook *et al.*, 2019). Interestingly, associative learning in mice also led to sparse population coding, where the total network activity was decreased while the response of cortical neurons was increased, presumably to provide efficient sensory processing of the learned stimuli (Gdalyahu *et al.*, 2012).

The economic allocation of neural resources may point to an interesting design evolved in animals with a compact neural network. Minimizing the number of neurons allocated for encoding, and thus maintaining a sparse coding strategy, may maximize memory capacity (Rolls and Treves, 1990). Furthermore, sensory neuroplasticity alters dynamic responses to various environmental cues, and consequently, innate behavioral outputs may be modulated. Thus, it may become advantageous to minimize the number of sensory neurons undergoing functional changes so that a larger fraction of the sensory system can preserve innate functions. In support of the minimal sensory units that can store a memory in *C. elegans*, the AFD neurons had been shown to store the recently experienced temperature in a cell-autonomous manner (Luo, Cook, *et al.*, 2014; Tsukada *et al.*, 2016). In addition, a few dedicated sensory neurons participate in learning and memory of salt concentrations (Luo, Wen, *et al.*, 2014; Jang *et al.*, 2019).

Integration of the distributed short-term memory information, encoded within individual sensory neurons, was observed in the immediate downstream layer of the interneurons (Fig. 3). We identified two interneurons, AIA and RIA, whose activity was modulated in a stimulus- and valence- specific manner, both following short- and long-term learning. The AIA interneurons are postsynaptic to most of the chemosensory neurons (White *et al.*, 1986; Cook *et al.*, 2019), and had been shown to integrate various chemosensory inputs (Shinkai *et al.*, 2011; Larsch *et al.*, 2015; Dobosiewicz, Liu and Bargmann, 2019). While AIA activity promotes forward locomotion (Garrity *et al.*, 2010), we found that long-term aversive training increased AIA activity together with increase in reversal rates (Fig. 3D-F and Fig. 4E-F). This may suggest that the AIA neurons are primarily integrating sensory modalities rather than directly affecting motor outputs.

In contrast, the RIA interneurons regulate head steering, and consequently movement directionality, by integrating perceived sensory cues with motor-neuron feedback that signals head position (Hendricks *et al.*, 2012; Hendricks and Zhang, 2013; Ouellette *et al.*, 2018). We found that these neurons also store and integrate past stimuli and valence experiences (Figure 3 G-L). Anatomically, the RIA neurons receive direct synaptic inputs from AWC and ASH, and indirect inputs from AWA (White *et al.*, 1986; Cook *et al.*, 2019), possibly explaining how these neurons may integrate memory information distributed within individual sensory neurons.

It is notable that LTAV-trained worms divert from the CS once approaching it (Fig. 4G-H). The altered choice of direction (suppl. Fig. S15H) and the neuronal activity in RIA (Fig. 3H) show correlating logic of regulation: both are markedly increased in LTAV. This might suggest that the modulated activity in the RIAs may underlie the change in movement directionality. In addition, RIA ablation mildly reduces reversal rates (Gray, Hill and Bargmann, 2005), raising the possibility that these neurons also contribute to the enhanced turning frequency and aversion in LTAV-trained worms (Fig. 4E-F).

Clearly, additional interneurons which control turning rates (e.g., AIB, AIZ, AIY, and RIM (Wakabayashi, Kitagawa and Shingai, 2004; Iino and Yoshida, 2009; Garrity *et al.*, 2010; Li *et al.*, 2014; Lee *et al.*, 2019), and speed (e.g., AIY, RIB, SIA, and RMG (Li *et al.*, 2014; Lee *et al.*, 2019)), may also be participating in these memory-induced behavioral modulations. Indeed, some of these neurons had been shown to participate in either memory formation (AIB & RIM) or memory retrieval (AIY & RIA (Jin, Pokala and Bargmann, 2016)). In addition, while we exclusively focused on neurons whose activity was modulated upon memory retrieval, these and other neurons may contribute to memory formation via transcriptional changes (Lakhina *et al.*, 2015; Freytag *et al.*, 2017). Together, interneurons might act as secondary memory cells that integrate and store information from the upstream primary sensory neurons. As activity of interneurons highly correlates with locomotion behavior (Kato *et al.*, 2015), these neurons directly dictate adaptive locomotion outputs upon encountering the CS.

While long-term associative memories were not evident in the sensory layer, they markedly modified activity of the downstream interneurons (Fig. 3). This may hint to another intriguing principle for coding memories in a compact neural network: Plasticity in the sensory neurons is likely to modulate sensory responses to various cues, possibly affecting innate behavioral outputs. For short-term memories, the modulated behavior will be brief, but for long-term memories, the impact on behavior will be long lasting. Thus, for long-term memories, it may be advantageous to ‘clear up’ information stored within sensory neurons, and to relegate it to deeper layers of the network, so that animals could quickly resume innate responses. This relegation of long-term memories to deeper layers can be viewed as analogous to the transfer process taking place in mammalian brains, where hippocampal short-term memories are moved for long-term storage in cortical areas (Rothschild, 2019). In addition, clearing information from sensory neurons may also mitigate the limited sensory resources as more neurons will become available for coding new short-lived memories.

Taken together, here we have systematically studied how associative memories are encoded within a compact neural network. The comprehensive cellular-resolution network-wide analysis revealed numerous principles that allow efficient encoding under the constraints of limited neural resources. These coding principles may well extend to memory formation and storage in higher organisms with more complex brain systems.

## Material and Methods

section is available in the supplementary information file.

## Data availability

All neuronal activity and behavioral data together with the associated analysis scripts are available in https://osf.io/5v4qu/.

## Acknowledgements

We thank Cornelia Bargmann and Einav Gross for strains. Some strains were provided by the CGC, which is funded by the NIH Office of Research Infrastructure Programs (P40 OD010440). This study was funded by ERC (336803), ICORE, and ISF (1300/17) to AZ. COP postdoctoral fellowship is supported by the David-Herzog-Funds at Styrian Universities. EB, RR, and EI are also supported by the Jerusalem Brain Center. AZ is the Joseph H. and Belle R. Braun Senior Lecture chair and the Greenfield chair in Neurobiology.

