## Supplementary information and figures for "Principles for coding associative memories in a compact neural network"

### Material and Methods

#### Worm cultures

Animals were grown at 20 °C on 9 cm nematode growth medium plates, seeded with 500 µL of confluent OP 50 bacterial suspension. For culturing and experiments, eggs were collected by dissolving the animals using standard bleaching protocols. The eggs were seeded at a density of 1000-1200 per plate. To inflict short-term memory, bleaching and seeding were conducted 3 days prior to the experiment. Animals undergoing long-term training were seeded 48 h before initiation of the training.

#### Training procedures

To induce olfactory associative memories, butanone was presented to the animals in combination with food (appetitive) or in the absence of food (starvation, aversive conditioning). Associated mock-trained control groups received the same treatment without butanone presentation. Naive control animals were non-treated worms of the same age.

**Short-term appetitive training.** Animals were first starved for one hour in 1 mL of M9 in a 15 mL centrifuge tube with an open lid. Worms were then trained on high-food NGM plates (seeded with 500 µL of confluent OP 50 culture) in the presence of 20 x 5 µL droplets of 10 % (v/v in DDW) butanone applied to the inside face of the plate lid. The mock-trained group received 20 x 5 µL droplets of DDW instead. Training duration was 1 h (modified after (Kauffman et al., 2010)).

**Short-term aversive training.** As described previously (Bargmann, Hartwieg and Robert Horvitz, 1993), worms were washed three times with an M9 buffer and transferred onto chemotaxis plates (no food). The trained group was incubated for 1 h with 20 x 5 µL droplets of 10 % (v/v in DDW) butanone, while the mock-trained group was incubated with an equivalent amount of DDW.

**Long-term appetitive training.** This training was performed by seven repetitions of starvation and food-butanone pairings as described in (Kauffman et al., 2010), with the addition of a mock-trained group. In contrast to all other training regimes, 5 x 2 µL droplets of 10 % (v/v) were used in long-term appetitive training because higher levels of butanone led to aversive choice behavior (data not shown).

**Long-term aversive training.** Worms were washed three times in an M9 buffer and transferred to chemotaxis plates. The trained group was starved with 20 x 5 µL droplets of 10 % (v/v) butanone on the lid for 10 h, with one exchange of the butanone droplets after 5 hours, while the mock-trained groups were starved in presence of DDW droplets.

### Behavioral assays

Behavioral changes induced following training were assessed using either two-choice (Fig. 1A), or one-choice (Fig. 4A) assays:

**Two-choice assays.** worms were washed three times with a chemotaxis buffer, and 100-200 animals were transferred onto the center point of the chemotaxis plate, 3.5 cm from the target endpoints. 4-pole and 2-pole layouts were used as depicted in suppl. Fig. S1A. Endpoints were loaded with butanone (butanone dissolved either in water or EtOH) or the alternative choice (diacetyl in water or pure ethanol, see suppl. Fig. S1A). Note that different concentrations of butanone were used for animals with appetitive and aversive training because of the valence-specific shift in choice behavior (see suppl. Fig. S2). Positively-trained animals were tested with 10-fold diluted butanone ($10^{-1}$) and negatively-trained animals were tested with 1000-fold diluted butanone ($10^{-3}$). Worms were immobilized at the endpoints by applying 1 µL of 1 M $NaN_3$. Animals in each region were subsequently scored to provide the choice index, and a learning index was calculated based on these choice indices as a measure of behavioral changes induced by learning (Kauffman *et al.*, 2011).

**One-choice assays.** Worms were washed three times with a chemotaxis buffer and transferred onto a chemotaxis plate. At a distance of 5 cm from the starting point, an agar plug with 3 µL of pure butanone was mounted to the inner face of the plate lid. Two cm from the starting point, we placed another agar plug supplemented with 3 µL of water. Animals were allowed to move freely for 30 minutes. Chemotaxis behavior was imaged using a Micropublisher 5 RTV CCD camera (QImaging, Canada) equipped with a ZOOM 7000 Navitar macro objective (Navitar, USA). Animal tracks were extracted using a multi-worm tracker (Itskovits *et al.*, 2017), from which we quantified accumulation rate at the endpoint, deviation angles, speed, and reversal rates. To control for butanone evaporation and to ensure behavior consistency, only the first 10 minutes of the movies were analyzed. To provide higher accuracy of local deviation angles and speed, tracks were segmented into 24-frames segments. The deviation angle is the angle between the vector pointing from the animal towards the endpoint and the average vector of the worm track segment (suppl. Fig. S15A,B). Due to the use of the center-mass-tracking, reversals were defined as any perceptible form of backward movement (Gray, Hill and Bargmann, 2005).

### Calcium imaging and data analyses

In preparation for live imaging and prior to loading the animals onto the microfluidic chips (Chronis, Zimmer and Bargmann, 2007), the animals were starved for 20 minutes on empty NGM plates. For imaging multiple neurons, worms were also paralyzed using 10 mM levamisole dissolved in chemotaxis medium. The worms were habituated to the restraint and paralysis for additional 10 minutes within the chip.

Neuronal activities were recorded for 90 seconds: 30 seconds after the initiation of imaging, animals were presented with 336 mM butanone (diluted in chemotaxis medium), and then imaging continued for additional 60 seconds. After the animals were acclimated to the presence of butanone for 5 minutes, we re-initiated imaging, and after 30 seconds of imaging we switched the stimulus (butanone) off, and continued imaging for additional 60 seconds.

A Nikon A1R+ confocal laser scanning microscope (Nikon, Japan) equipped with a 40x 1.15 NA water immersion objective was used for fast live imaging. Z-series of the head region of the animal was recorded at 0.9-2 volumes per second depending. Individual Z-stacks were scanned at 0.4-0.8 μm intervals in the sensory-reporter line (ZAS280) and the pan neuronal reporter line (ZAS319). For imaging neurites of single AIA and RIA neurons, worms were not paralyzed. The stimulus exchange interval was 20 seconds and responses were recorded for 3 minutes without interruption. Z-Stacks were acquired at 0.5 - 0.8 μm intervals. The z-stack sampling rate was 2-5 Hz. Single sensory neurons were imaged with an IX 83 epifluorescence microscope (Olympus, Japan) and a 40x 0.95 NA objective. The acquisition was controlled by μManager (Edelstein *et al.*, 2010).

To identify individual neurons in multi-neuron z-stack time series, neuronal somas were segmented using a Gaussian fitting and a tracking algorithm (Toyoshima et al., 2016) targeting nuclear mCherry tags. GCaMP intensities in target neurons were extracted from segmented neurons by a custom-built analysis pipeline in Matlab (Mathworks, USA) reading voxels within a 70 percent radius of the initial segmentation radius. Image stacks from neurites were projected by summing all images using imageJ. Projected micrographs were analyzed using custom imageJ and Matlab scripts utilizing Fiji's trackMate plugin (Schindelin et al., 2012). Since data were acquired with varying frame rates, neuronal activation graphs were linearly interpolated to a 2 Hz sampling rate for multi-neuron datasets and a 5 Hz for the neurite datasets.

Neural activation levels were normalized by their ground state ($F/F_G$), unless stated otherwise.

For sensory neurons, the ground state was extracted from the last 10 frames (after visual inspection) of the imaging, long past the stimulus switch, and after the neuron resumed its ground pre-exposure state. This was particularly important for neurons like ASH and ASK which strongly responded to the blue imaging light showing markedly increased activities prior to butanone addition/removal. For interneurons, a 10-frame ground state was visually identified. This is because interneurons often stochastically transition between active and inactive states which precludes extracting ground states using a fixed time window. Notably, similar results were obtained when normalizing by the 10-frames time window prior to the switch (data not shown); however, the use of ground-state levels for normalization was more accurate. For statistical comparisons, intensities following stimulus exchange were summed. Integration times were neuron-specific since neuronal dynamics strongly varied among neurons. For AWA, AWC, AWB, RIAnrD, RIANrV, and AIA neurons 10 seconds; RIAnrS, 12 seconds; ASH, 15 seconds; AVA, 20 seconds; ASJ and ASI, 30 seconds.

### Statistical analysis

Statistical analysis of neuronal activation and behavior was done in MatLab and Python. Data were tested for normal distribution by Shapiro-Wilk test (small sample number) or Kolmogorov-Smirnov test (larger sample number). ANOVA or ANOVA on ranks followed by pairwise comparisons based on t-tests or Wilcoxon rank-sum test/signed rank test, depending on the underlying distribution, were used to test for differences. Multiple comparisons were adjusted using false discovery rates, FDR (Benjamini and Hochberg, 1995). In the neurite dataset, neuronal intensity measurements within a 95% confidence interval were used. For comparing neuronal activation between the different learning paradigms, neurons with reliable activity responses were included in the analysis (namely, AWA, ASH, $AWC^{ON}$, $AWC^{OFF}$, AWB, ASJ, ASI, AVA, RIAnrD, RIAnrV, RIAnrS, and AIA). For each of these neurons, twelve pairwise comparisons of activity post stimulus exchange were conducted as stated in table 1, yielding 156 comparisons in total. The same comparison matrix was also used for locomotion parameters presented in Figure 4.

Significant experience-dependent changes in choice behavior were detected by one-sample t-tests against the zero value. Comparing learning indices of mutants or ablated animals with wildtype controls was done by pairwise comparisons.

**Table 1. pairwise comparisons of the different conditions.**

| Comparisons design | |
|---|---|
| **Group 1** | **Group 2** |
| STAP-T | STAV-T |
| LTAP-T | LTAV-T |
| STAP-T | LTAP-T |
| STAV-T | LTAV-T |
| STAP-T | STAP-M |
| STAV-T | STAV-M |
| LTAP-T | LTAP-M |
| LTAV-T | LTAV-M |
| STAP-T | NAIVE |
| STAV-T | NAIVE |
| LTAP-T | NAIVE |
| LTAV-T | NAIVE |

STAP, Short-term appetitive. STAV, Short-term aversive. LTAP, Long-term appetitive, LTAV, Long-term aversive. T-Trained. M-Mock.

### Worm strains

For functional imaging, worm strains driving expression of calcium reporters in neurons of interest were used. Behavioral analysis experiments were conducted with wild type worms, mutants, and genetic neuronal ablation lines as listed in suppl. table 2.

**Supplementary Table 2. Worm strains used in this study.**

| Designation | Phenotype/ Purpose | Genotype/Expression (Source/Ref) |
|---|---|---|
| **Behavioral analysis:** | | |
| N2 | wild type | N2 wild type (CGC) |
| PY7502 | AWC ablation | [P*ceh*-36::TU#813, P*ceh*-36::TU#814, P*srtx-1*::GFP, P*unc-122*::DsRed] (Beverly, Anbil and Sengupta, 2011) |
| CX4 | AWA functional loss | *odr*-7(ky4) (Sengupta, Chou and Bargmann, 1996) |
| JN1713 | ASH ablation | [P*sra*-6::mCasp-1 + P*unc*-122p::mCherry] (Yoshida *et al.*, 2012) |
| **Functional imaging:** | | |
| CX16561 | AIA reporter line | *[*P*gcy28d*::GCaMP D381Y coel::dsRed, P*odr*-7::Chrimson::SL2::mCherry,P*elt*-2::mCherry 2] (Larsch *et al.*, 2015) |
| P*gcy*-37::YX2.60 | URX reporter line | [P*gcy*-37::YX2.60] (Gross *et al.*, 2014) |
| PS6250 | CEPD reporter line | [P*dat*-1::GCaMP3] (Zaslaver *et al.*, 2015) |
| PS6374 | $AWC^{ON}$ reporter line | [P*str*-2::GCaMP3] (Zaslaver *et al.*, 2015) |
| PS6253 | $AWC^{OFF}$ reporter line | [P*srsx*-3::GCaMP3] (Zaslaver *et al.*, 2015) |
| ZAS97 | $AWC^{OFF}$ and $AWC^{ON}$ reporter line | [P*str*-2::ChR2-cherry,P*str*-2::GCaMP3,*srsx*-3::GCaMP3] - This work |
| ZAS96 | $AWC^{ON}$ reporter line in *unc-13* bkg | [P*str*-2::GCaMP3] in *unc-13(e51*) - This work |
| ZAS76 | $AWC^{ON}$ reporter line in *unc-31* bkg | [P*str-2*::GCaMP3] in *unc-31*(*e928*) - This work |
| ZAS280 | Sensory neuron reporter | [P*osm*-6::GCaMP3, P*osm*-6::ceNLS-mCherry-2xNLS] (Iwanir *et al.*, 2019) |
| ZAS319 | pan-neuronal reporter line | AML-32 (Nguyen *et al.*, 2017) x ZAS280 [P*rab*-3::NLS::GCaMP6s + P*rab-3*::NLS::tagRFP, P*osm*-6::GCaMP3, P*osm*-6::ceNLS-mCherry-2xNLS] - This work |
| ZAS42 | RIA reporter line | [P*glr-3*::GCaMP3] - This work |

### Identifying individual neurons

Neurons in the sensory neurons reporter line (P*osm-6*::GCaMP3) and in the pan-neuronal reporter line (P*rab*-3::NLS::GCaMP6s) were identified by comparing with anatomic maps (White *et al.*, 1986; Durbin, 1987) in a custom-built Matlab 3D visualization tool. Neurons that could not be unambiguously determined, were verified by comparing anatomic position and activity profiles based on GCaMP reporter lines of individual identified neurons. This approach was used for identifying $AWC^{ON}$, $AWC^{OFF}$, and URX neurons (see suppl. Fig. S6&S10).

We used the *osm-6*::GCaMP line (ZAS280) from which we could unambiguously identify virtually all chemosensory neurons (of both lateral sides), except for the AFD neurons for which the signal was too dim to provide reliable measurements. Since $AWC^{ON}$/$AWC^{OFF}$ neurons randomly appear on either the right or the left side of the nerve ring, we used an additional reporter line that discriminates between these cell fates (see below). Also, since we could not discriminate between the URX and CEPD neurons due to their proximity, we used two additional reporter strains, each expressing a calcium indicator exclusively in one of these neurons (see below).

**Identifying and discriminating between the $AWC^{ON}$ and the $AWC^{OFF}$ neurons.** The AWC class of neurons consists of two neurons, $AWC^{ON}$ and $AWC^{OFF}$, where the $AWC^{ON}$ neuron exclusively expresses the gene *str-2*, and the $AWC^{OFF}$ exclusively expresses the gene *srsx-3* (Wes and Bargmann, 2001; Zaslaver *et al.*, 2015). However, the bilateral position of each of these neurons (that is, right or left) is stochastically determined (Sagasti *et al.*, 2001).

When using the P*osm-6*:GCaMP line, we found that one neuron (could be the right- or the left-side neuron in the different animals) exhibited strong responses to butanone in naïve, trained- and mock-trained animals, while its bi-lateral neuron showed weak responses in naive and mock-trained animals, but significantly stronger responses following short-term training paradigms (Fig. 2B, D and suppl. Fig. S10). However, using the P*osm-6*::GCaMP line to simultaneously image all chemosensory neurons on both lateral sides of the worm, did not allow assigning which one is $AWC^{ON}$ and which is $AWC^{OFF}$.

To differentiate between the two neurons and assign each with its corresponding activity, we made use of additional strains in which only one the two AWC neurons expresses GCaMP (PS6253, PS6374, Table 2), and a strain where the two neurons are differentially tagged: both AWC neurons express GCaMP using the *str-2* ($AWC^{ON}$) and *srsx-3* ($AWC^{OFF}$) promoters, $AWC^{ON}$ also expresses a red tag via P*str-2*::ChR2-mCherry (ZAS97, Table 2).

We repeated the training and imaging experiments with these strains to find that $AWC^{OFF}$ response dynamics to butanone is strong and robust in both, trained- and mock-trained animals. In contrast, $AWC^{ON}$ showed negligible responses in the mock-trained animals but its activity was significantly elevated following short-term training paradigms, (both positive and negative, suppl. Fig. S10).

**Discriminating between the URX and CEPD neurons.** The two neuron types URX and CEPD are located in close proximity in the dorsal part of the nerve ring. When using the P*osm-6*::GCaMP line (ZAS280) to simultaneously measure activity from all chemosensory neurons, we observed an activity in these neurons but could not assign it to one of these neurons. To identify and discriminate between the two neurons, we made use of two additional lines, each expressing a calcium reporter in only one of these neurons using a neuron specific promoter. These lines are: P*gcy-37*::YC2.60, exclusively expressed in URX (Gross *et al.*, 2014), and P*dat-1*::GCaMP (Zaslaver *et al.*, 2015), exclusively expressed in dopaminergic neurons of which CEPD is one of them (Zaslaver *et al.*, 2015).

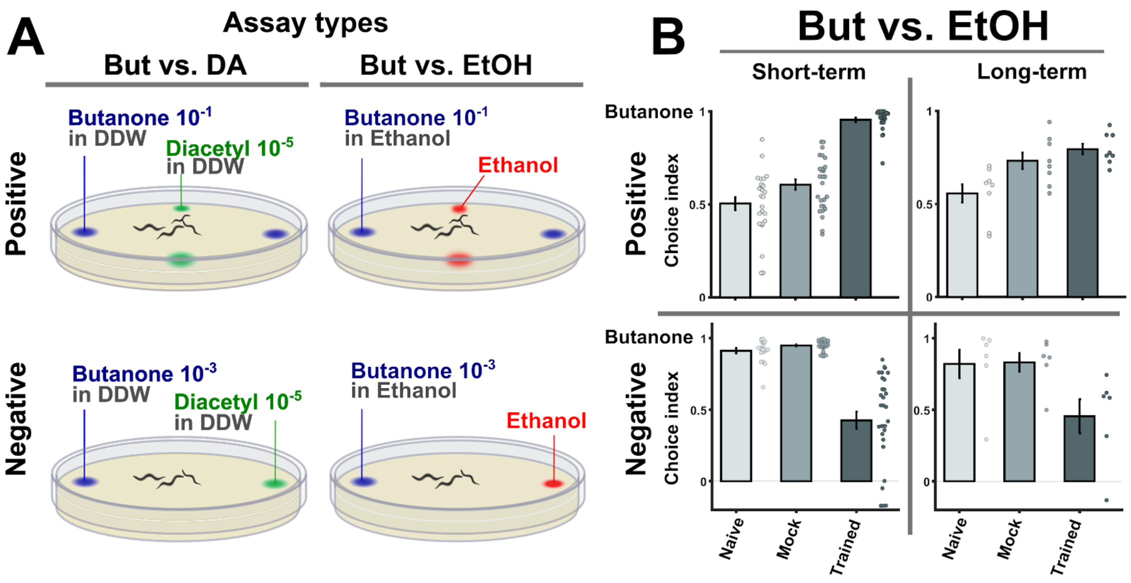


**Supplementary figure S1. Behavioral choice assays using ethanol as the alternative choice reproduced the results obtained using diacetyl as the alternative choice.**

**(A)** The layout of the 2-choice assay. Based on published methods (Bargmann, Hartwieg and Robert Horvitz, 1993; Kauffman *et al.*, 2010) a four quadrant layout was used for scoring the preference of animals that underwent positive associative training. A two opposing choices layout was used to score preference of animals that underwent negative (starvation) associative learning. Note that butanone concentrations were different for appetitive and aversive assays because the valence of training causes valence-specific shifts in the choice behavior (see suppl. Fig. S2). We performed these 2-choice assays for butanone vs. diacetyl and butanone vs. ethanol, which yielded comparable results.

**(B)** When using ethanol instead of diacetyl as the alternative choice (as shown in Fig. 1C), similar preference values to the CS butanone are obtained: Positively-trained animals are more attracted to butanone while negatively-trained animals are less attracted to it. This underscores the findings that the four training paradigms form robust memory traces that can be behaviorally observed.

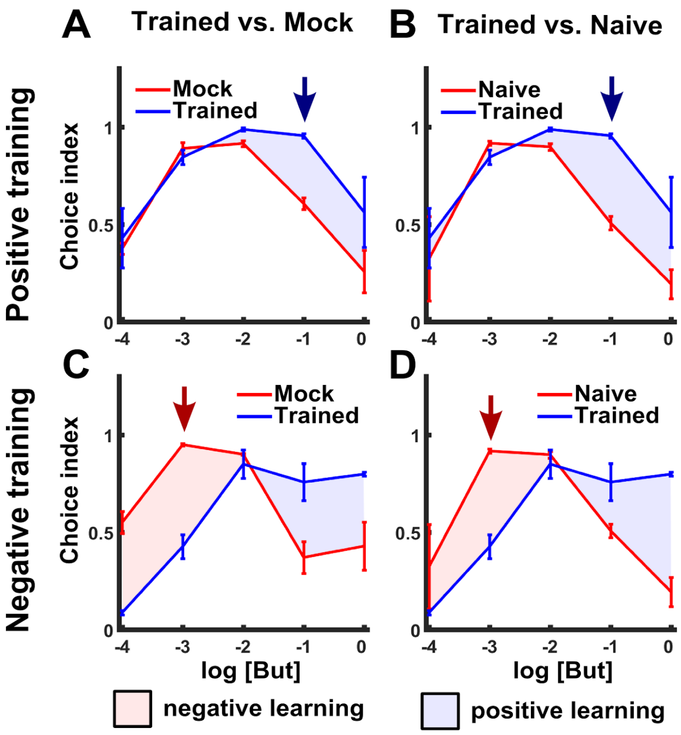


**Supplementary figure S2. Appetitive and aversive training paradigms shift the preference to butanone.**

**(A-B)** Positive training increased attraction towards higher concentrations of butanone (blue areas) when compared to mock-trained (A) or naive (B) animals over a range of butanone concentrations. We therefore used a concentration dilution of $10^{-1}$ (blue arrow) for the positive training (with food) and for the subsequent choice assays (Fig. 1 and suppl. Fig. S1).

**(C-D)** In contrast, following negative (starvation) training, animals' attraction towards butanone decreased, but only to the lower concentrations of butanone (compared to mock-trained controls (C) and naive animals (D), red areas). The maximal reduction was noted at the $10^{-3}$ dilution (red arrow) so we used this dilution in the choice assays (Fig. 1 and suppl. Fig. S1). For each concentration, we included 2-10 independent repeats, each consisting of 3 plates with ~100 animals. Error bars indicate SEM.

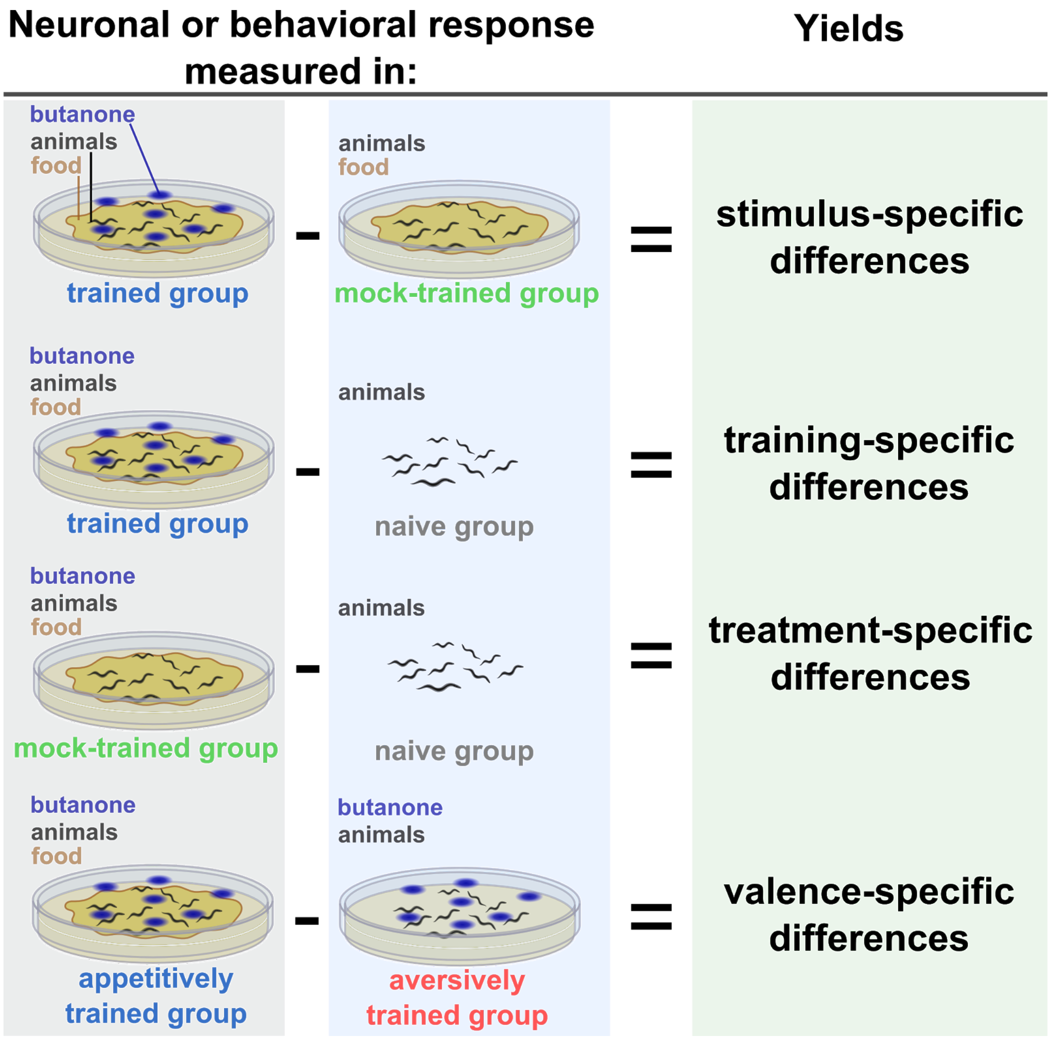


**Supplementary figure S3. Comparing performances (neural activity or behavior) of the different assayed/control groups allows inferring the cause that underlies the modulation.**

In each experiment (neuronal imaging or behavioral choice assays), in addition to the trained animals, we always included analysis of naive and mock-trained animals. Mock-trained animals underwent the same training treatment (e.g. starvation) but were unexposed to the conditioned stimulus (CS) butanone. Naive animals were left untreated but were assayed in parallel to the mock- and trained-animals. When comparing behavioral preferences of the trained groups (aversive or appetitive) and the associated mock-trained controls, the impact of the CS (smell of butanone) becomes evident. Therefore, the observed differences are termed **stimulus-specific**. When trained animals are compared to naive animals, differences introduced by the entire training procedure (US+CS) become apparent. These differences are referred to as **training-specific**. To control for the changes introduced merely by the treatment itself (e.g. food availability), we compared behavioral outputs between mock-trained and naive animals (termed **treatment-specific)**. When comparing negative training to positive training, the differences are due to the context of the experience (starvation or food availability), and are hence termed **valence-specific**. Notably, due to the high day-to-day variability in behavioral outputs, it was mandatory to compare between the various control groups within the same experimental day (see Fig. 1D and suppl. Fig. S4). Thus, we used the different control groups to derive the learning indices (LIs) as shown in Fig. 1B & D.

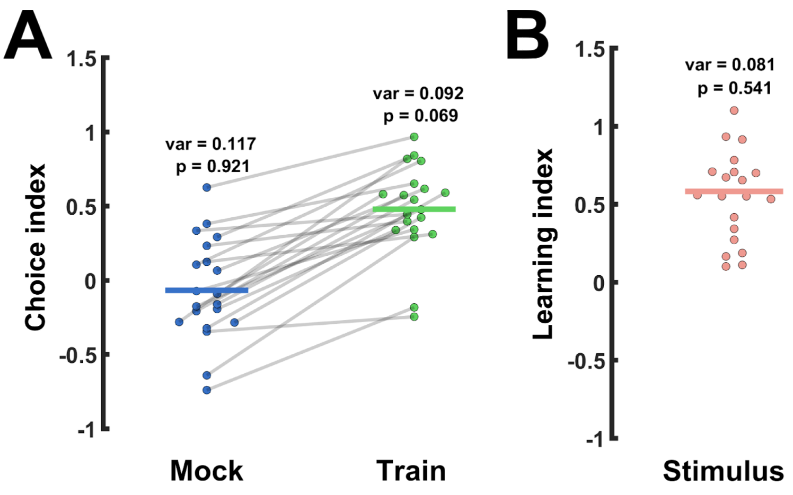


**Supplementary figure S4. Behavioral choice assays are subject to high day-to-day variability.** Variability can be reduced by comparing same-day choice indices (CIs). This was done by calculating learning indices (LIs, see Fig. 1B,D) which denoted CI differences among the various groups that were assayed on the same day.

**(A)** CI values widely vary within each tested group (e.g. shown are short-term positively-trained and mock-trained groups as provided in Fig. 1C). Note that lines connect two groups assayed on the same day. Thus, when considering same-day results, the significance becomes apparent as trained animals always enhanced attraction towards the CS (positive training).

**(B)** When expressing the change in the CIs as the stimulus-specific learning index ($CI_{trained}$ – $CI_{mock\text{-}trained}$), the variance decreases (12.0% less than in $CI_{trained}$ and 30.8% less than in $CI_{mock\text{-}trained}$). Measurements are likely to stem from a normal distribution (high p-value in Shapiro-Wilk test, p). As a consequence, testing Learning Indices against 0 presents an option independent from naive day-to-day variability. Variance = var, p = p-value of a Shapiro-Wilk test. Horizontal lines indicate the mean.

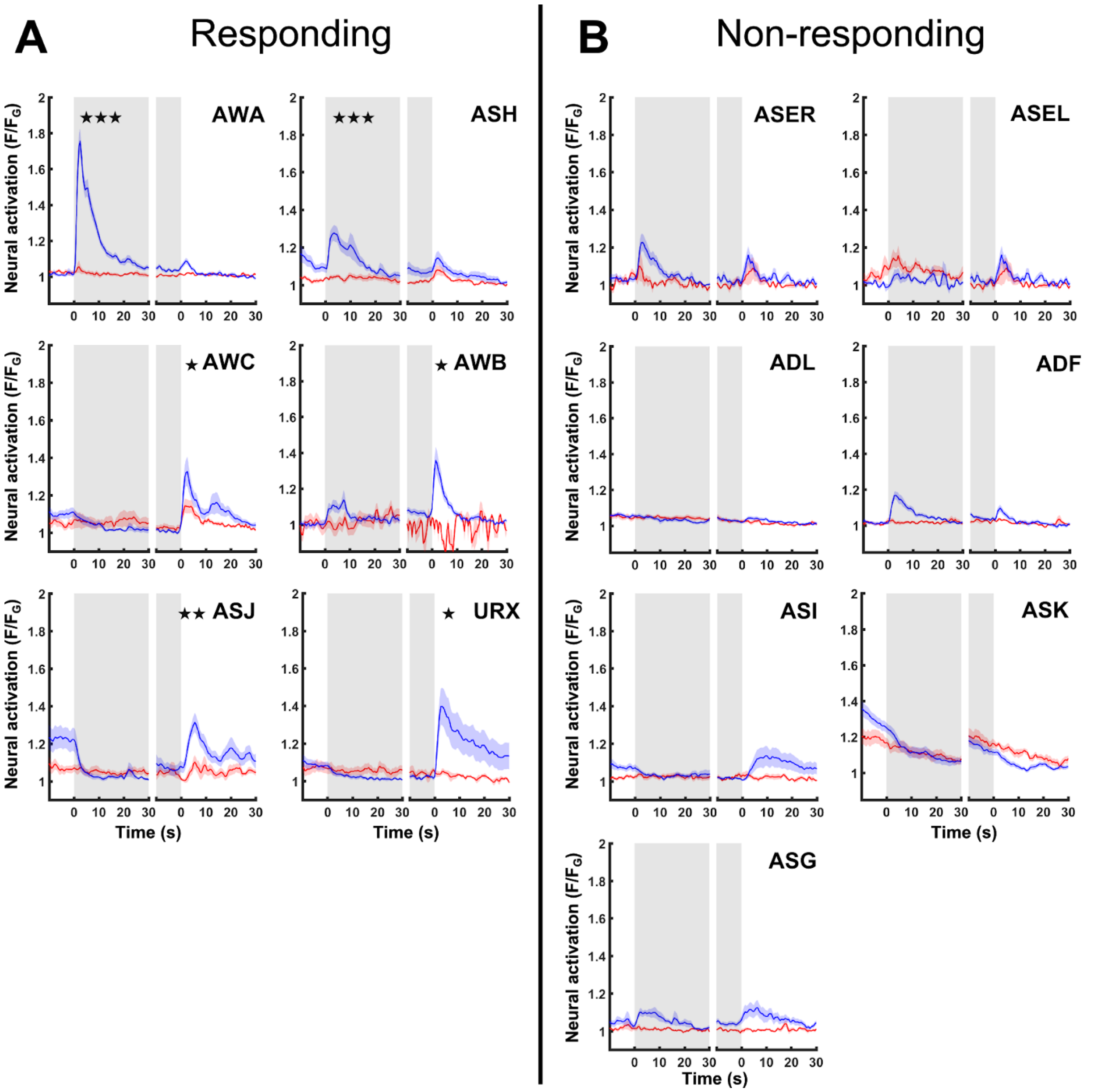


**Supplementary figure S5. Elucidating the chemosensory neurons that show innate responses to butanone.**

**(A)** The set of chemosensory sensory neurons that show innate responses to butanone presentation or removal. * $p<0.05$, ** $p<0.01$, *** $p<0.001$(t-test, FDR corrected).

**(B)** The set of chemosensory neurons that did not show significant responses to butanone presentation or removal.

Notably, in all imaging experiments, we added rhodamine to the butanone solution to accurately time butanone presentation or removal following ON and OFF switches. To account for possible evoked responses due to rhodamine itself (which included chloride ions in its solution) we measured neural responses following exchange of buffers where only rhodamine was added to only one of the buffer streams. In all panels, blue curves denote responses to butanone+rhodamine (presentation or removal). Red curves serve as controls and denote responses to buffer exchanges lacking butanone but supplemented with rhodamine solution. Gray and white areas denote presence or absence of butanone, respectively (for the blue curves) and presence or absence of rhodamine, respectively (for the red curves).

AWA, ASH, AWC, AWB, ASJ, and URX neurons showed robust responses to butanone exchange in naive animals (compared to the buffer controls). However, URX was not included in further analysis since we could not reliably read from this neuron throughout all conditions. ASI neurons did not show significant increased activations compared to buffer controls. However, this neuron was included in analysis because it exhibited differential activation throughout the different conditions (see suppl. Fig. S7). In the set of non-responding chemosensory neurons, ADF and ASG showed weak activity but since they are located in close proximity to the AWA neurons, which exhibited strong activations, we could not exclude possible bias due to cross-reads. The ASE neurons showed low responses to rhodamine only, probably due to the chloride ions in the rhodamine solution. These responses were comparable to the butanone responses, hence, we categorized them as non-responding to butanone. Since the response dynamics of the right and left symmetry neurons was similar (except for the ASE neurons), we present their averaged response dynamics.

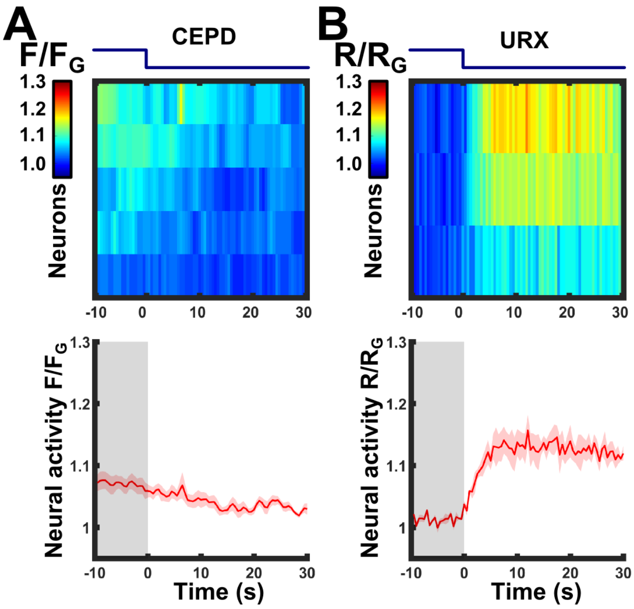


**Supplementary figure S6. Butanone removal elicits responses in the URX neurons, but not in the CEPD neurons.** Using the *osm-6*::GCaMP strain (Fig. 2A), we observed a response to butanone removal in the dorsal side of the lateral ganglion. Since both the CEPD and the URX neurons are located in close proximity in that region, it was impossible to distinguish between them and tell which neuron is actually responding. We therefore used strains expressing calcium reporters exclusively in either of these neurons. These experiments showed that the URX neurons respond to butanone removal.

**(A)** CEPD neurons do not respond to butanone off-step. Imaging the strain *dat-1*::GCaMP (PS6250 (Zaslaver *et al.*, 2015)) which drives expression in the dopaminergic neurons and which CEPD is one of them. N=5 animals.

**(B)** Neural responses in a strain expressing cameleon in URX (P*gcy-37*::YC2.60 (Gross *et al.*, 2014)). A clear sharp response is detected in the URX neurons upon butanone removal (N=3 animals). The blue line above the heat maps indicates the time course of the stimulus switch (on to off); Bottom panels, mean responses of the traces shown in the upper panels. Shaded-red areas around the mean curves indicate SEM.

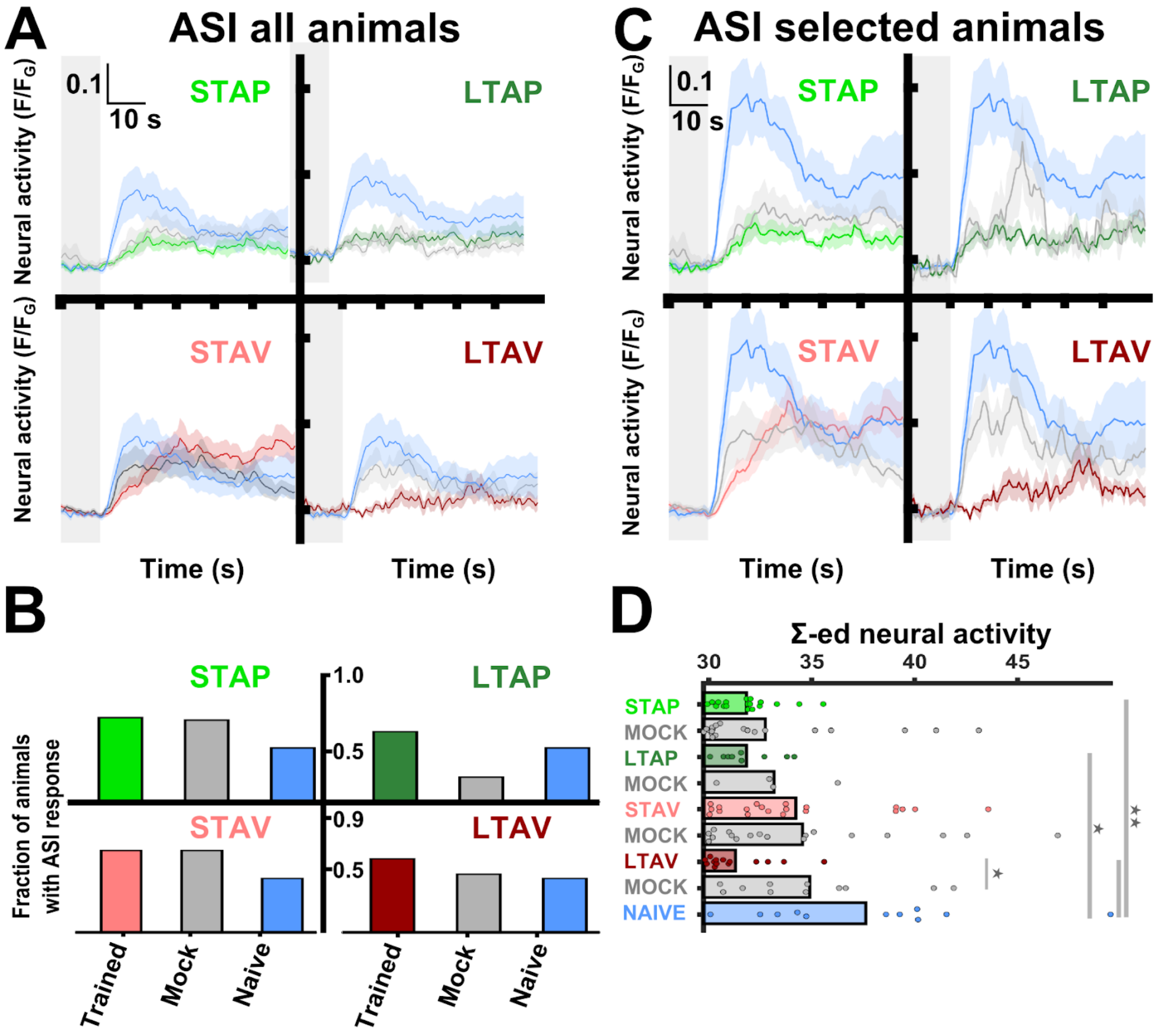


**Supplementary figure S7. Response dynamics of the ASI neurons are modulated following memory formation.**

**(A)** Activity dynamics of the ASI neurons across the four training paradigms. The shaded gray background indicates presence of butanone. The number of worms tested to compute the mean response for the different conditions and control groups ranges between 7-12. Shaded-colors of the curves denote SEM. Shaded gray rectangles indicate presence of butanone (Same for panel C).

**(B)** Fraction of animals in which the ASI neurons responded. ASI responses were observed in only ~half of the animals. As this population heterogeneity was particularly evident for the ASI neurons, we averaged ASI activity based on the fraction of responding animals (panel C). Bars indicate the portion of animals exceeding the 13% activation threshold post-stimulus removal. This threshold was selected based on the background noise of the neural activity extracted from control experiments, and which was 8.9±4.2% (see suppl. Fig. 5).

**(C)** Activity dynamics of ASI neurons that responded above the 13% threshold. Notation as for panel A.

**(D)** When considering only animals in which the ASI neurons responded above the threshold, activity of trained and mock-trained animals is significantly reduced across all training paradigms. ASI responses were modulated following positive training in both short-and long-term association paradigms. Furthermore, activity STAV training is significantly higher compared to STAP training, possibly indicating a valence-specific coding. *p<0.05, **p<0.01, ***p<0.001 (t-test, FDR corrected). Error bars are SEM.

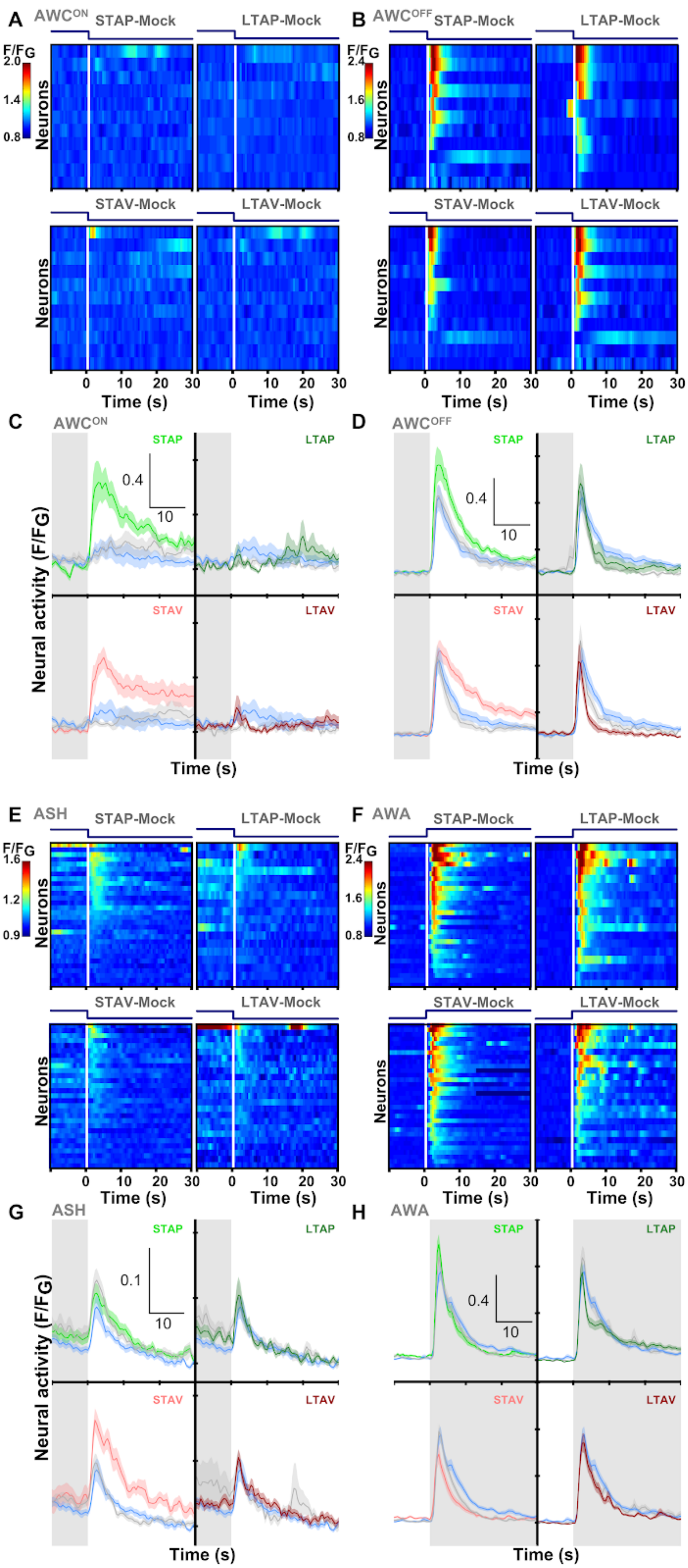

A
AWC^ON
STAP-Mock
LTAP-Mock
STAV-Mock
LTAV-Mock
F/FG
2.0
1.4
0.8
Neurons
Time (s)
B
AWC^OFF
2.4
1.6
C
D
STAP
LTAP
STAV
LTAV
0.4
10
Neural activity (F/FG)
E
ASH
1.6
1.2
0.9
F
AWA
G
0.1
H

**Supplementary figure S8. Short term, but not long-term, memories are evident at the sensory layer.**

The sensory neurons, $AWC^{ON}$ (A,C), $AWC^{OFF}$ (B,D), ASH (E,G), and AWA (F,H) are modulated following short-term training paradigms.

Shown are the individual neural response traces of mock-trained animals and below are their mean neural activities (gray) along with activities of naive animals (blue) and of trained animals (color coded by the training paradigm). Shaded gray background indicates butanone presence. Shaded areas in the line plots indicate SEM.

**(A,C) Activity of $AWC^{ON}$**. (A) Heat maps of individual traces of mock-trained animals. (C) The $AWC^{ON}$ neuron becomes responsive following short-term training paradigms only: aversive (STAV, red) and appetitive (STAP, light green). No differences were observed between naive, mock-trained and long-term appetitive (LTAP, dark green) or long-term aversive (LTAV, dark red) trained animals. Neuronal activities in STAV and STAP are significantly higher than in the associated mock controls, suggesting that this neuron codes for the stimulus (see Figure 2B for p-values) .

**(B,D) Activity of $AWC^{OFF}$**. (B) Heat maps of individual traces of mock-trained animals. (D) Activity of $AWC^{OFF}$ is enhanced following short-term training paradigms. Both STAV- and STAP-trained animals exhibit significantly increased responses when compared to naive or the corresponding mock-trained animals (see Figure 2C for p-values).

**(E,G) Activity of ASH neurons**. (E) Heat maps of individual traces of mock-trained animals. (G) Activity of the ASH neurons increases following STAV training only. This enhanced activity is significantly higher than the matched mock-trained control animals and the trained LTAV animals, suggesting that the ASH neurons may code for the stimulus and the valence (see Figure 2D for p-values).

**(F,H) Activity of AWA neurons**. (F) Heat maps of individual traces of mock-trained animals. (H) Activity of the AWA neurons is dampened following STAV training only. It is significantly lower than activity of STAP-trained and naive animals but not different from associated mock-controls, indicating that AWA neurons may code for the valence (see Figure 2E for p-values).

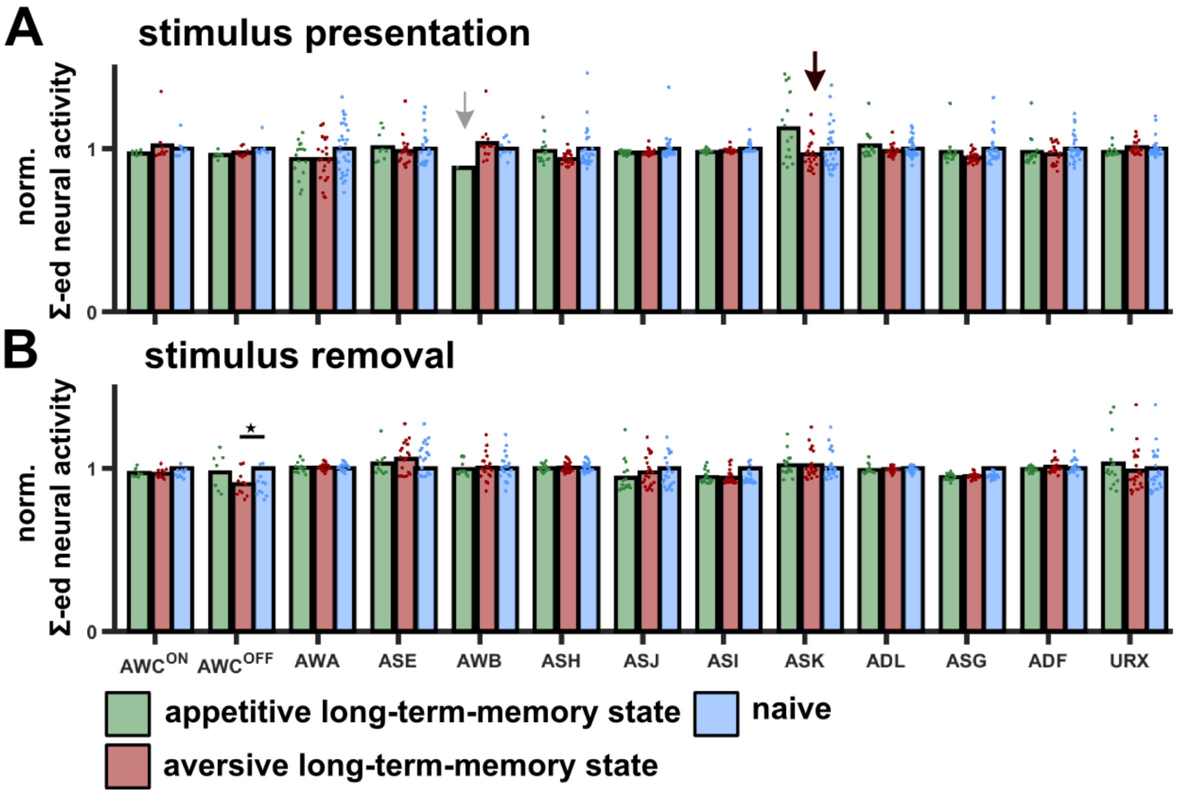


**Supplementary figure S9. Responses of chemosensory neurons following long-term training were not different from the responses observed in naive animals.**

Normalized summed neuronal activities following butanone presentation **(A)** or removal **(B).** These summed responses are normalized by each neurons' naive response. The AWC$^{OFF}$ neuron showed a minor (yet significant) reduced activity following LTAV training. *$p < 0.05$ (t-tests, FDR corrected). Note that for the AWB neurons (gray arrow), we did not collect enough neuronal activations to perform a statistical comparison for the stimulus presentation. However, AWB neurons are not activated by butanone on-steps. ASK neurons (black arrow) show a significantly heightened activity in response to the blue light that was used for imaging. These responses are not related to butanone presentation.

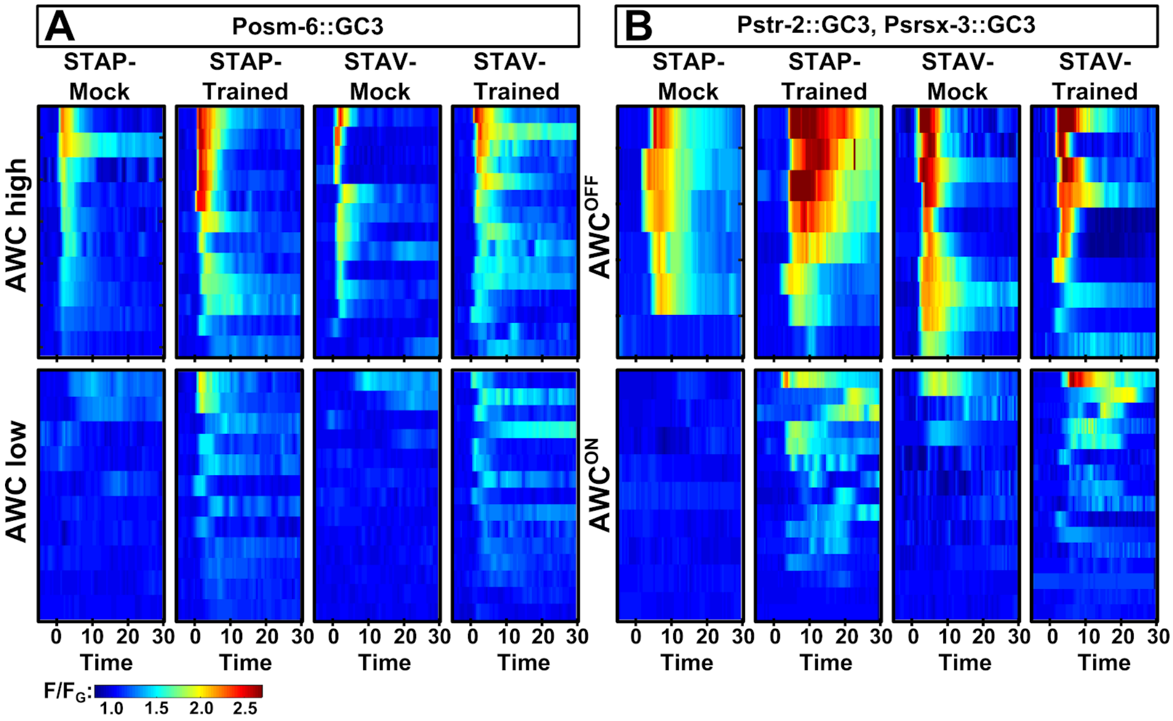


**Supplementary figure S10. Discriminating between the two AWC neurons.**

**(A)** When imaging the activity of both AWC neurons using the ‘all-chemosensory’ P*osm-6*::GCaMP line, we found that the two AWC neurons exhibited distinct responses. One neuron showed high responses to butanone (denoted as AWC high) in trained and mock-trained animals for both short-term positive (STAP) and short-term aversive (STAV) training paradigms. The other neuron exhibited weak responses to butanone in mock-trained animals (denoted as AWC low), but robust strong responses to butanone following STAV and STAP.

**(B)** Imaging AWC neuronal activity in strains where AWC-subclass identity is known. In these lines, $AWC^{OFF}$ shows strong responses to butanone in both trained and mock-trained animals. These dynamics correspond to the AWC high neuron shown in panel A. Accordingly, $AWC^{ON}$ matches the dynamics corresponding to AWC low. In both panels, butanone was removed at time point 0 (off-step response).

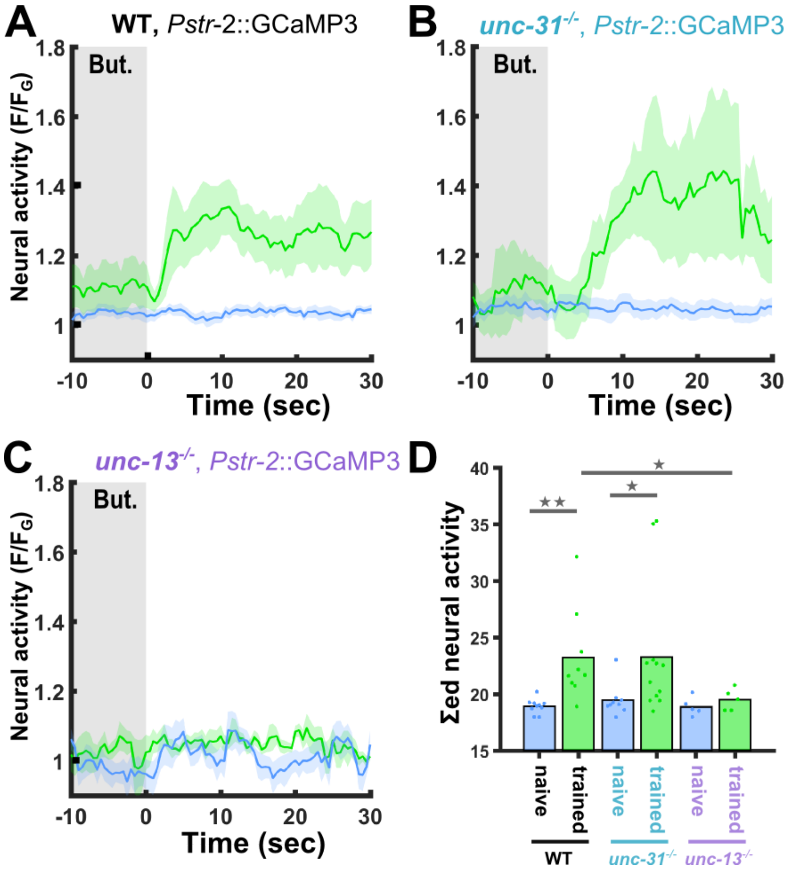


**Supplementary figure S11. The gained responses in AWC$^{ON}$ following short-term appetitive training requires intact synaptic transmission.**

**(A)** In naive wildtype animals, the AWC$^{ON}$ neuron does not respond to butanone removal (blue). Short-term positive training modulates activity of the AWC$^{ON}$ neuron which becomes responsive to butanone (blue, naive: 10 animals; green, trained: 9 animals).

**(B)** In animals defective in neuropeptide release (*unc-31*), AWC$^{ON}$ dynamics is similar to the one observed in wild type animals as shown in panel A. (blue, naive: 9 animal; green, trained 12 animals).

**(C)** In animals defective in synaptic transmission (*unc-13*), the increased response in the AWC$^{ON}$ was abolished (blue, naive: 5 animals; green, trained: 5 animals).

**(D)** Bar graphs showing summed neuronal activity 15 seconds following the stimulus exchange. Blue naive animals, green trained animals. Dots in the bar graph indicate repeats. Shaded areas in line plots and indicate SEM. *$p<0.05$, **$p<0.01$ (t-test, FDR corrected).

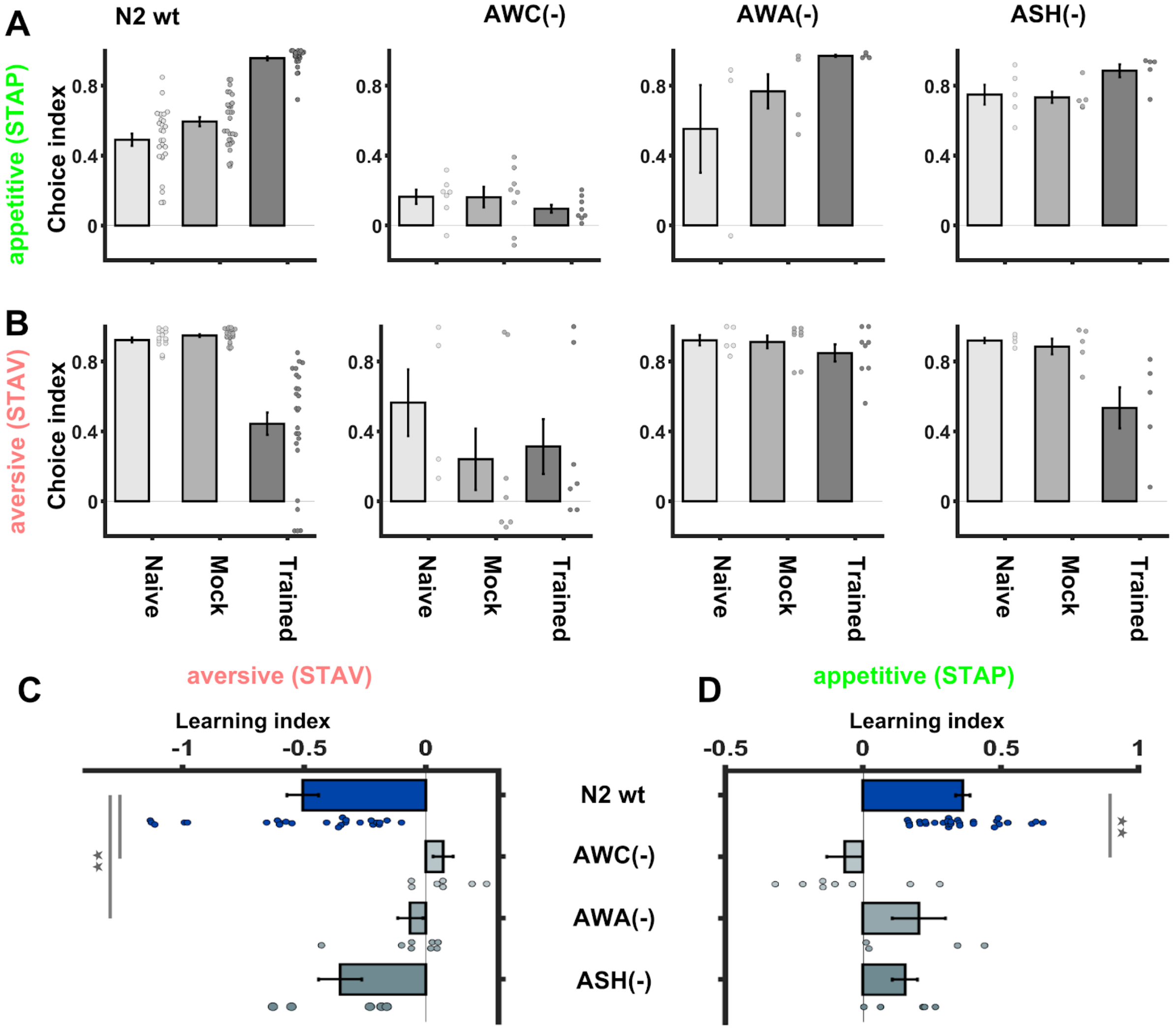


**Supplementary figure S12. Lack of functional AWC or AWA neurons impairs formation of short-term memories.**

**(A-B)** Choice Indices (CIs) of WT and mutant strains following short-term appetitive (A) and short-term aversive (B) training. AWC(-) (Beverly, Anbil and Sengupta, 2011) and ASH(-) (Yoshida *et al.*, 2012) are genetically-ablated strains. AWA(-) are mutants with dysfunctional AWA neurons (*odr-7*$^{-/-}$). Animals were assayed using the two-choice assay with the CS butanone and EtOH as the alternative choice (as shown in suppl. Fig. S1).

**(C-D)** Stimulus-specific learning indices ($CI_{Trained}$ - $CI_{Mock\text{-}trained}$), as shown in Fig. 1B, were calculated based on the CI values shown in (A & B). Animals lacking AWC neurons and animals with dysfunctional AWA neurons showed learning deficits following STAV- and STAP-training paradigms. Values are the means of 4-8 independent experimental repeats for each strain (29 for WT N2). Error bars denote SEM. **$p<0.01$ (rank-sum test, FDR corrected).

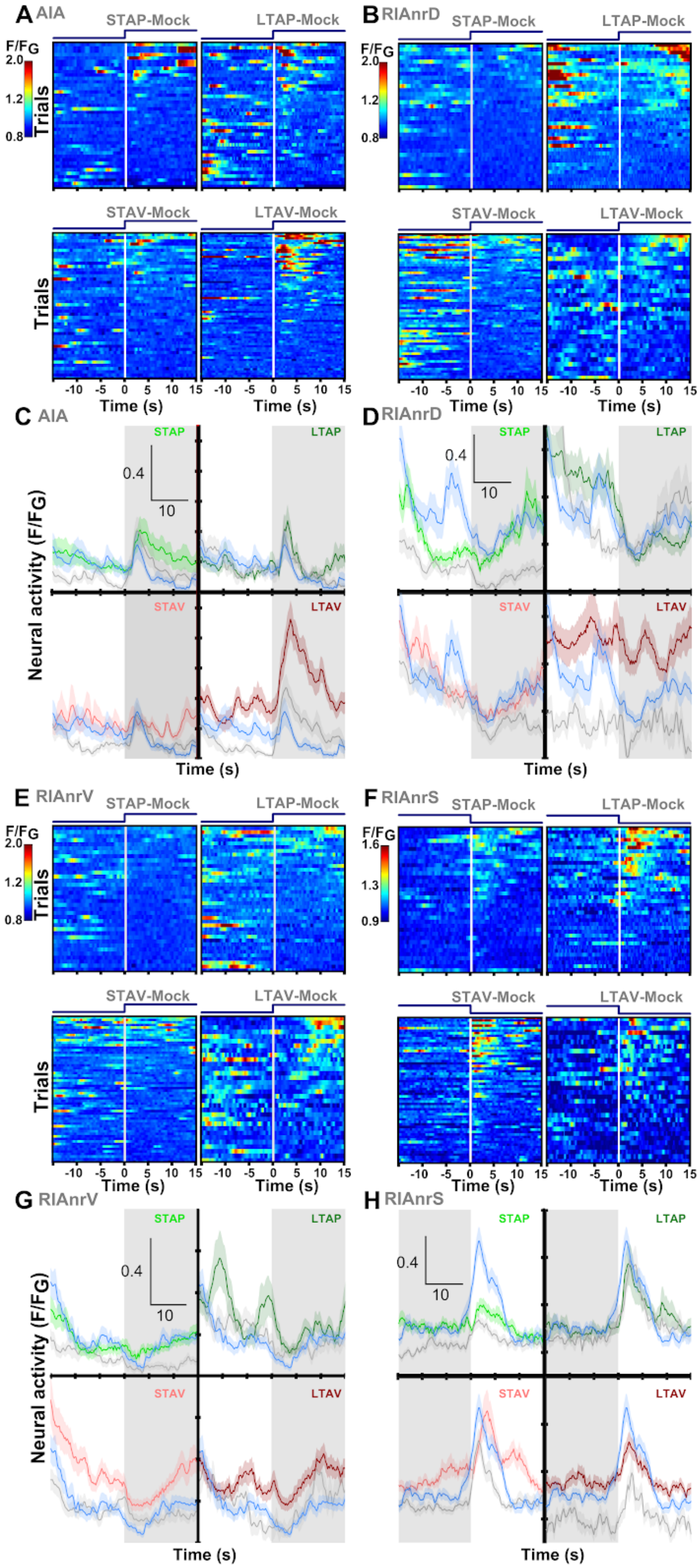
A AIA
STAP-Mock
LTAP-Mock
F/FG
2.0
1.2
0.8
Trials
STAV-Mock
LTAV-Mock
Trials
-10 -5 0 5 10 15
Time (s)
B RIAnrD
STAP-Mock
LTAP-Mock
F/FG
2.0
1.2
0.8
STAV-Mock
LTAV-Mock
Time (s)
C AIA
STAP
LTAP
0.4
10
Neural activity (F/FG)
STAV
LTAV
Time (s)
D RIAnrD
STAP
LTAP
0.4
10
STAV
LTAV
Time (s)
E RIAnrV
STAP-Mock
LTAP-Mock
F/FG
2.0
1.2
0.8
Trials
STAV-Mock
LTAV-Mock
Trials
Time (s)
F RIAnrS
STAP-Mock
LTAP-Mock
F/FG
1.6
1.3
0.9
STAV-Mock
LTAV-Mock
Time (s)
G RIAnrV
STAP
LTAP
0.4
10
Neural activity (F/FG)
STAV
LTAV
Time (s)
H RIAnrS
STAP
LTAP
0.4
10
STAV
LTAV
Time (s)

**Supplementary figure S13**. **The interneurons AIA and RIA show modulated activity that is stimulus- and valence-specific.** This figure supplements Fig. 3 in the main text which shows activity of trained animals. Here we show the raw data for the mock-trained groups.

**(A,C)** Activity traces extracted from AIA neurites. (A) heat-maps of individual trials in mock-trained animals. (C) Population means of AIA activity in animals following training in the different paradigms with the corresponding mock-trained and naive animals. Following LTAV training, activity in AIA neurites was significantly higher compared to STAV and mock-controls (see p-values in FIg. 3) (C). Note, LTAV mock-trained controls also show an increased activity, though significantly lower.

**(B,D)** Activity traces extracted from the dorsal neurites of the RIA neurons (RIAnrD). (B) Heat-maps of individual trials in mock-trained animals. (D) Population means of activation. Note that in LTAV-trained and mock-trained animals there is a substantial difference in baseline activity before stimulus presentation and in the event-related activity after butanone presentation (for p-values see Figure 3).

**(E,F)** Activity traces extracted from the ventral neurites of the RIA neurons (RIAnrV). (E) Heat-maps of individual trials in mock-trained animals. (F) Population means of neural activation.

**(G,H)** Activity traces representing sensory-evoked signals in RIA neurites (RIAnrS). (G) Heat-maps of individual trials in mock-trained animals. (H) Population means of activity in animals trained in the different paradigms. The response to stimulus removal is significantly higher in STAV-trained animals compared to associated mock-controls. Moreover, it is higher in LTAV animals compared to their associated mock-controls, suggesting that this neuron integrates both stimulus and valence information (p-Values see Figure 3).

In (C,D,G, H): Blue, naive; Gray, mock trained; Color coded, trained animals. Shaded gray areas indicate presence of butanone. Shaded areas in the line plots indicate standard error of the mean.

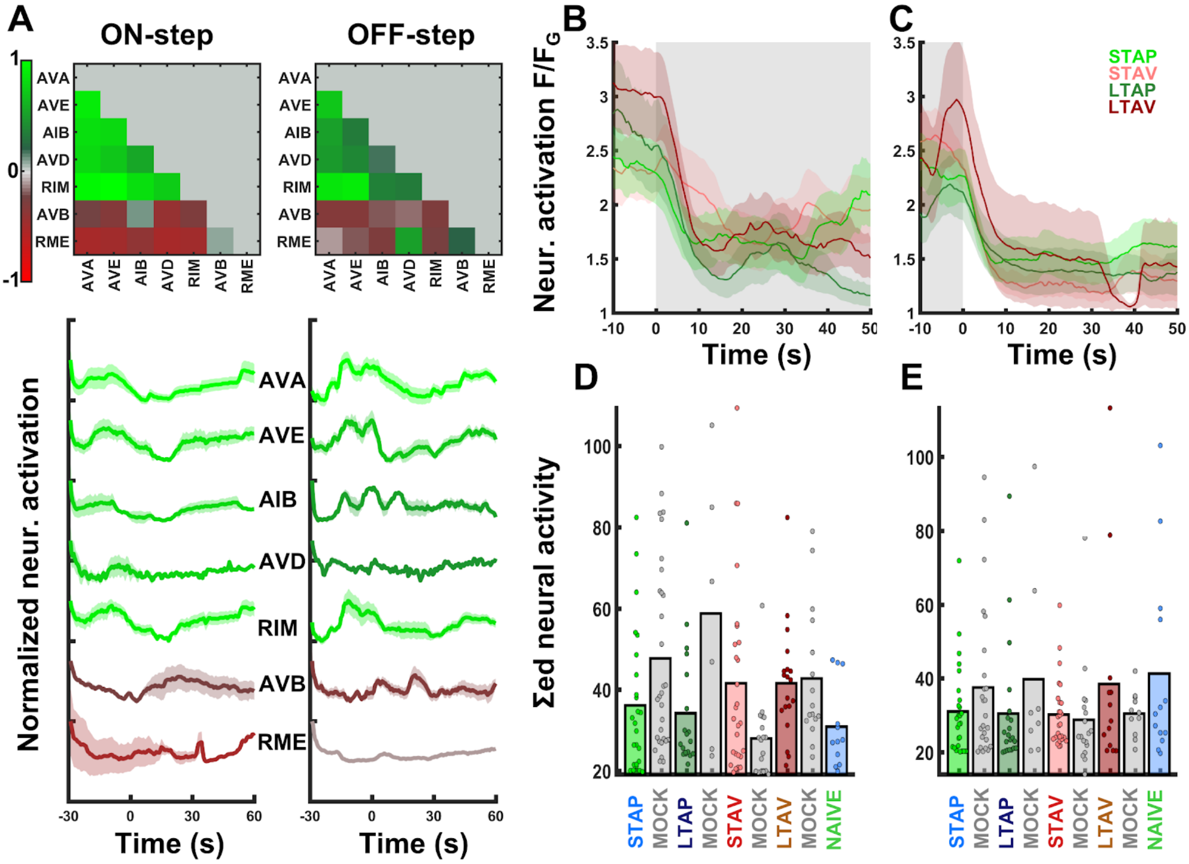


**Supplementary figure S14. Activity of command neurons was not modulated following training in the four different paradigms.**

**(A)** Average pairwise Pearson correlation coefficients of activity extracted from the soma of command neurons and other interneurons (of naive animals). The activity profiles below denote mean neuronal activity of each neuron. Green and red activity profiles show neurons that are highly correlated with AVA activity. Red profiles, neurons with anti-correlated activity. Error interval shades denote SEM. The command neurons oscillated between activated and inactivated states in intervals on the order of tens of seconds. In fact, as animals were imaged while being restrained and paralyzed, activity of command and interneurons presumably reflected fictive locomotion that deviated from the natural state of the animal (Scholz *et al.*, no date; Kato *et al.*, 2015). As AVA neurons could be segmented with the highest reliability, and since its activity was highly correlated (anti-correlated) with other command neurons, we used the AVA activity profile as a non-redundant indicator for the activity of all other command neurons.

**(B-C)** Mean activity profiles of the AVA neurons following exposure to (A) or removal of (B) the CS butanone in each of the four training paradigms. While AVA neurons responded to stimulus exchange by switching from active to inactive states, there were no significant differences between the conditions following the stimulus exchange.

**(D-E)** Comparing the summed neural activity 20 second post stimulus exchange yields no significant differences upon presentation (C) or removal (D) of butanone. Notably, other integration times showed similar results (data not shown).

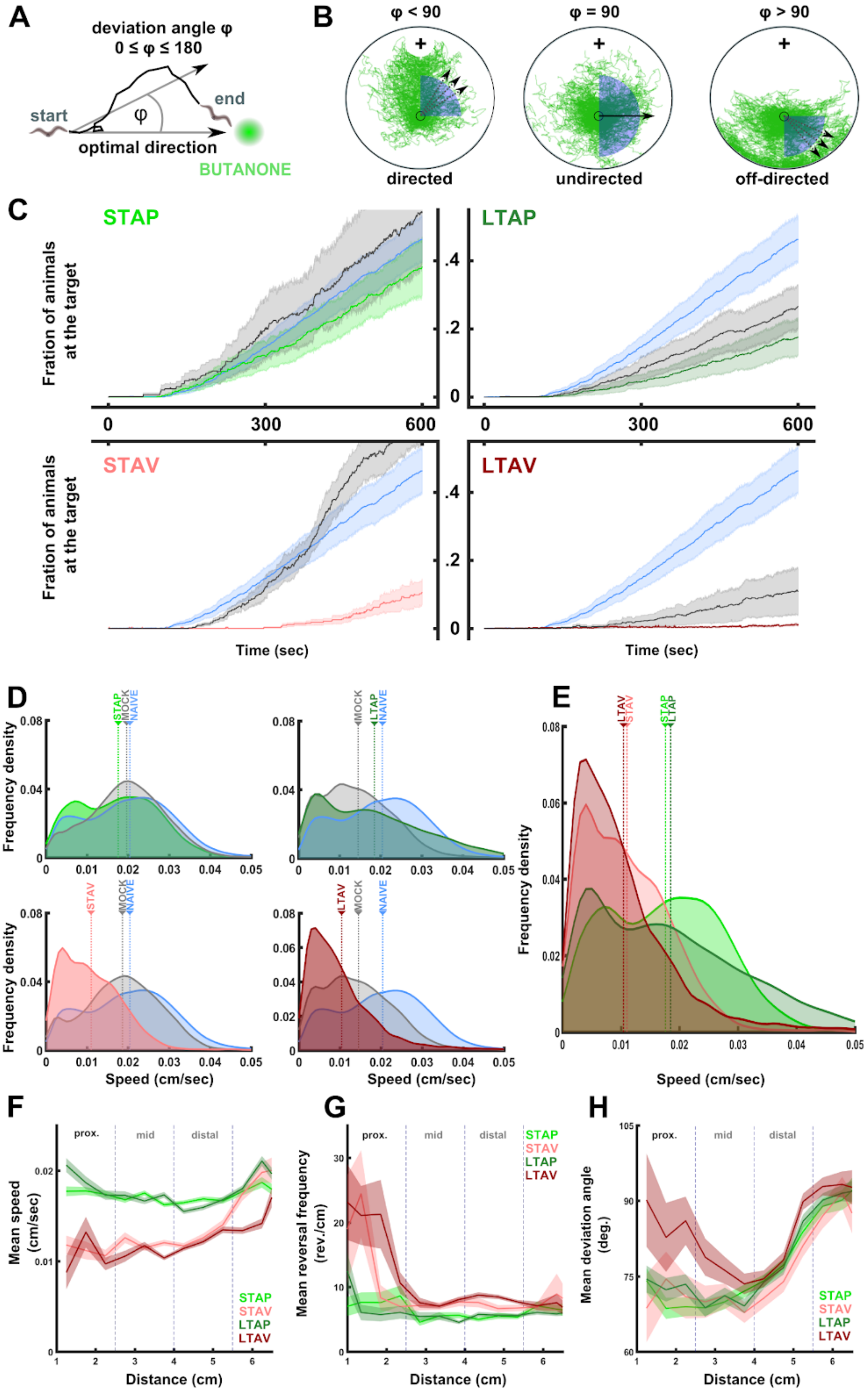
A
deviation angle φ
0 ≤ φ ≤ 180
start
end
φ
optimal direction
BUTANONE
B
φ < 90
φ = 90
φ > 90
directed
undirected
off-directed
C
STAP
LTAP
STAV
LTAV
Fration of animals at the target
Time (sec)
D
E
STAP
STAV
LTAP
LTAV
MOCK
NAIVE
Frequency density
Speed (cm/sec)
F
G
H
prox.
mid
distal
Mean speed (cm/sec)
Mean reversal frequency (rev./cm)
Mean deviation angle (deg.)
Distance (cm)

**Supplementary figure S15. Comparing deviation angles, speeds and reversal rates during chemotaxis following training across all four paradigms.**

**(A)** Deviation angle is calculated from 24-frame track segments and defined as the angle between the optimal direct vector to the end target and the average movement vector during the segment interval. Resulting angles range between 0-180 degrees.

**(B)** Tracks with deviation angles lower than 90° are directed towards the chemoattractant end point (+). Tracks with a deviation angle at 90° are undirected. Tracks with a deviation angle greater than 90° are facing away from the chemoattractant.

**(C)** Accumulation of animals reaching the target CS over time for each of the training paradigms. Blue, naive. Color coded in each quadrant are the specific training paradigms. Gray, corresponding mock-trained animals. Notably, we found substantial differences between the training paradigms in the dispersal timing of animals at the beginning of the assay. Naive animals and animals trained via the short-term paradigms started to disperse and chemotax almost immediately after they were placed in the start area. In contrast, animals trained through the long-term paradigms, including the associated long-term mock controls, exhibited a markedly slower dispersal as they remained in piles at the initiation site and left the starting point at later time points (visually detected). This difference is probably caused by treatment-specific effects (e.g. habituation to touch as a result of the training protocol) and indeed is evident in both long-term trained and mock animals. This is the reason why accumulation of these animals, particularly the positively-trained animals, lags behind and remains overall low. However, the Metrics shown in D-H are not affected by this slower dispersal.

**(D)** Locomotion speed distributions of track segments proximal to the chemoattractant for trained animals, associated mock-trained animals and naive animals. **(E)** Locomotion speed distributions proximal to the chemoattractant for all trained conditions. The speed of aversively-trained animals is lower than the speed of positively-trained animals (see Fig. 4D for p-values)

**(F)** Overlay of locomotion speed across all conditions. Note the decrease in locomotion speed of aversively-trained animals (see Fig 4 D for p-values).

**(G)** Reversal rates of trained animals (colored according to the paradigm). Aversively-trained animals show a significant increase in reversal rates upon approaching the CS target (proximal region, p-values see Fig. 4 F).

**(H)** Deviation angles of trained animals (colored according to the paradigm). Note that LTAV is the only condition markedly increasing deviation angles in proximity of the stimulus. (p-values are in Fig. 4 H).
